# A GABARAP−PtdIns3K-C1 positive feedback loop at the heart of the phagophore nucleation

**DOI:** 10.64898/2026.03.17.712327

**Authors:** Antoine N. Dessus, Yohei Ohashi, Maxime Bourguet, Tomos E. Morgan, Anastasia Nunez, Maria Manifava, Nicholas T. Ktistakis, Roger L. Williams

## Abstract

Macroautophagy/autophagy is a cellular process enabling degradation of intracellular components during starvation. In mammalian cells, autophagosomes can reach diameters of over 1000 nm within 30 min after triggering starvation, but how such substantial amounts of membranes can be synthesized within a brief time remains elusive. A protein complex central to the phagophore initiation is the lipid kinase PIK3C3-Complex 1 (PtdIns3K-C1), which produces phosphatidylinositol-3-phosphate (PtdIns3P). PtdIns3P recruits a variety of downstream proteins, among which is PtdIns3P-binding WIPI2 that facilitates lipidation of mammalian ATG8 (mATG8) family proteins on phagophores. Here we show that upon inhibition of mATG8 lipidation in cells, there is a decreased accumulation of WIPI2, suggesting a feedback loop between mATG8s and PtdIns3P production. The role of PtdIns3K-C1 in this feedback was demonstrated by in vitro experiments where recombinant membrane-coupled mATG8s bind to and potently activate PtdIns3K-C1, with GABARAP being the most potent activator among all mATG8s. By a combination of cryo-electron microscopy, structural mass spectrometry, activity assays and mutagenesis, we show that GABARAP binds two sites in PtdIns3K-C1, with one site showing an atypical bipartite interaction with the mATG8. We also confirm both sites are essential for GABARAP to activate PtdIns3K-C1. We propose that once GABARAP is indirectly recruited by PtdIns3P generated by basal activity of PtdIns3K-C1, a positive feedback loop is formed where PtdIns3K-C1 interacts with GABARAP and becomes activated to produce more PtdIns3P, thereby further stimulating GABARAP lipidation. This mechanism would be central for autophagosome biogenesis, where enlarged membranes need to be synthesized within a brief period.

**Graphical abstract:** The GABARAP−PtdIns3K-C1 positive feedback loop.
Model for the GABARAP−PtdIns3K-C1 positive feedback loop. GABARAP is indirectly recruited to the growing phagophore by PtdIns3P and activates PtdIns3K-C1, leading to an increased PtdIns3P production. The E1 (ATG7), E2 (ATG3) and E3 (ATG5-ATG12–ATG16L1) enzymes and WIPI2 are involved in the lipidation (covalent coupling) of GABARAP to membranes.

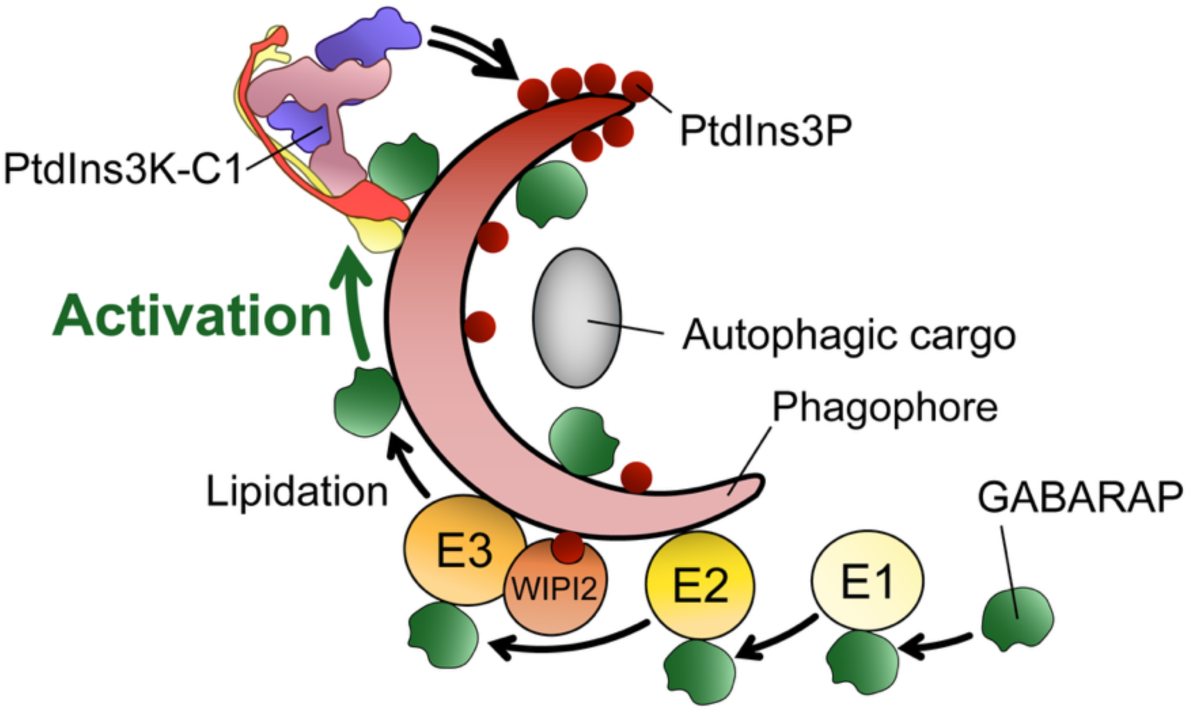

## Introduction

Autophagy defines pathways where cytoplasmic material too large to be degraded by the proteasome, like protein aggregates and organelles, are targeted instead to the lysosomes for degradation. This includes microautophagy, chaperone-mediated autophagy and macroautophagy (hereafter referred to as autophagy)^1–3^, and defects in autophagy have been linked to pathologies including cancer, neurodegenerative diseases, and inflammatory and immune disorders^4–7^. Biogenesis of autophagosomes involves nucleation, expansion, closure of membranes (phagophores or isolation membranes) that sequester cargos, followed by their fusion with lysosomes^8^. Autophagy is initiated at the endoplasmic reticulum (ER) where stress signals are integrated by the unc-51 like kinase complex ULK1, which recruits the class III PI3K complex 1 (hereafter referred to as PtdIns3K-C1)^9,10^. The human PtdIns3K-C1 is composed of the PIK3C3 lipid kinase, the PIK3R4 pseudokinase, ATG14 (BAKOR) and BECN1 proteins^11^. PIK3C3 and PIK3R4 constitute the kinase arm of the complex whereas BECN1 and ATG14 form its adaptor arm, giving a characteristic V-shape to PtdIns3K-C1^12–14^. PtdIns3K-C1 phosphorylates phosphatidylinositol (PtdIns) to yield phosphatidylinositol-3-phosphate (PtdIns3P), at a low basal activity that can be increased by the small GTPase RAB1A, which recruits PtdIns3K-C1 to membranes and activates it^15^.

PtdIns3P is essential for autophagosome generation because it recruits its effector WIPI proteins^16^. WIPI2d is known to interact with PtdIns3K-C1, and this interaction is believed to produce positive feedback^17^. Although it was shown that WIPI2d on PtdIns3P-containing membranes enhances the recruitment of PtdIns3K-C1 to membranes^17^, it remains unclear whether this results in an increase in the PtdIns3K-C1 activity. WIPI2d then recruits the ATG5-ATG12–ATG16L1 complex to autophagosomal membranes^18^. This complex acts as an E3 ligase involved in a ubiquitin-like enzymatic cascade along with the E1 ATG7 and the E2 ATG3. The substrates for this enzymatic cascade are Atg8 (yeast) or the mammalian-ATG8 proteins (mATG8s), which include LC3A, LC3B, LC3C, GABARAP, GABARAPL1 and GABARAPL2^19,20^. The Atg8/mATG8s become covalently coupled to the phagophore’s lipids by the E3 complex, allowing the recruitment of various receptors, which in turn recruit the autophagic cargo to be degraded^21^. The mATG8s thus bind to a plethora of proteins, which insert their LC3-interacting regions (LIRs) into the LIR-docking sites (LDSs) of the mATG8s. The LIR is a four amino acid sequence conserved among mATG8s binders, consisting of the motif Θ-X-X-Γ (were Θ is an aromatic residue, X any residue and Γ a small aliphatic residue) interacting with hydrophobic pockets in the LDS^22^. Recently, such LIR-LDS interactions were found in peptides from three subunits of PtdIns3K-C1: PIK3C3, BECN1 and ATG14, with a particularly strong affinity for the mATG8 GABARAP^23^. However, no binding of mATG8s to the full PtdIns3K-C1 assembly was attempted, and the functional implications on PtdIns3P production remained unknown.

Remarkably, during autophagy in mammalian cells, within a brief time^24,25^, one cell generates around 100 autophagosomes^26^, each 500–1500 nm in size^27^. To achieve this, a large amount of lipids is necessary, and the membranes need to be rapidly expanded and closed to engulf cargos and deliver them to lysosomes. Atg8/mATG8 proteins have roles in regulating autophagosome size^28^, elongation of phagophore membranes^29,30^, and autophagosome closure^31^. To execute these autophagic events efficiently, it is conceivable that many Atg8/mATG8 proteins need to be lipidated on the membranes in a brief time.

Here we show that the inhibition of mATG8 lipidation decreases the accumulation of the PtdIns3P effector WIPI2 relative to FIP200 in the cell, suggesting a reduction of PtdIns3P synthesis by PtdIns3K-C1. Using a combination of in vitro activity assays, cryo-electron microscopy (cryo-EM), structural mass spectrometry (MS), and mutagenesis we confirmed that lipidated mATG8s activate PtdIns3K-C1, and we identified two binding sites for the mATG8 GABARAP on PtdIns3K-C1. These two sites, namely the N site and the C site, both contribute to the activation of PtdIns3K-C1 by lipidated GABARAP on membranes. Our results suggest a positive feedback loop happens in cells between GABARAP and PtdIns3K-C1, which could be critical for rapid PtdIns3P production and associated mATG8 lipidation on the growing phagophore.

## Results

### Translocation of PtdIns3P effectors is sensitive to lipidation of mATG8s

We first examined the effect of the lack of mATG8 lipidation on its upstream PtdIns3P synthesis in the cell. A previous study showed that even in rich media, PtdIns3P-binding proteins WIPI1 and DFCP1 accumulated in MEF cells lacking the Atg5 gene that encodes an E3 subunit of the mATG8 lipidation machinery^32^. These results could have arisen from indirect effects over a long period of time in the Atg5 knockout cells. Because autophagy occurs in a brief time, we wanted to observe the acute effect of inhibition of mATG8 lipidation activity. To achieve this, we used an inhibitor of ATG7 which completely eliminates the lipidation of all mATG8 family members^33^, preventing their translocation to punctate structures upon autophagy induction (Fig. 1a, shown for LC3). Using this inhibitor, we determined whether translocation of early autophagy proteins to early autophagy structures was affected by the absence of lipidated mATG8, using as relevant markers FIP200 (a member of the ULK1 complex, which translocates first to autophagy puncta) and WIPI2 (an immediate effector of PtdIns3P which requires activation of PtdIns3K-C1 for its translocation to autophagy puncta). The number of FIP200 and WIPI2 puncta upon induction of autophagy with PP242 in the presence or absence of ATG7 activity was similar (Fig. 1b and c), suggesting the formation of the pre-autophagosomal structures is not affected by the loss of mATG8 lipidation. Interestingly, the intensity of FIP200 puncta was increased upon ATG7 inhibition (Fig. 1b small inserts and d), indicating an accumulation of FIP200 induced by stopping the autophagy flux downstream. However, the intensity of WIPI2 puncta was not increased upon ATG7 inhibition. This result contrasts to the WIPI accumulation seen in cells when the Atg8 lipidation is permanently deleted^32^, or when the autophagy flux is stopped downstream of mATG8s^18,34^. This suggests that the initial translocation of the early autophagy machinery to autophagy structures, which takes place upstream of mATG8 engagement, can occur in the absence of lipidated mATG8. However, the continuous translocation of the PtdIns3P effectors seemed to be affected by a transient loss of lipidated mATG8s, as WIPI2 does not further accumulate. Considering these results, and a previously characterised interaction between peptides derived from PtdIns3K-C1 subunits and mATG8s^23^, we hypothesised that the lipidated mATG8s may directly affect the PtdIns3P production by PtdIns3K-C1 in a feedback loop. We therefore carried out the in vitro characterisation of the interaction between these proteins.

**Figure 1.**
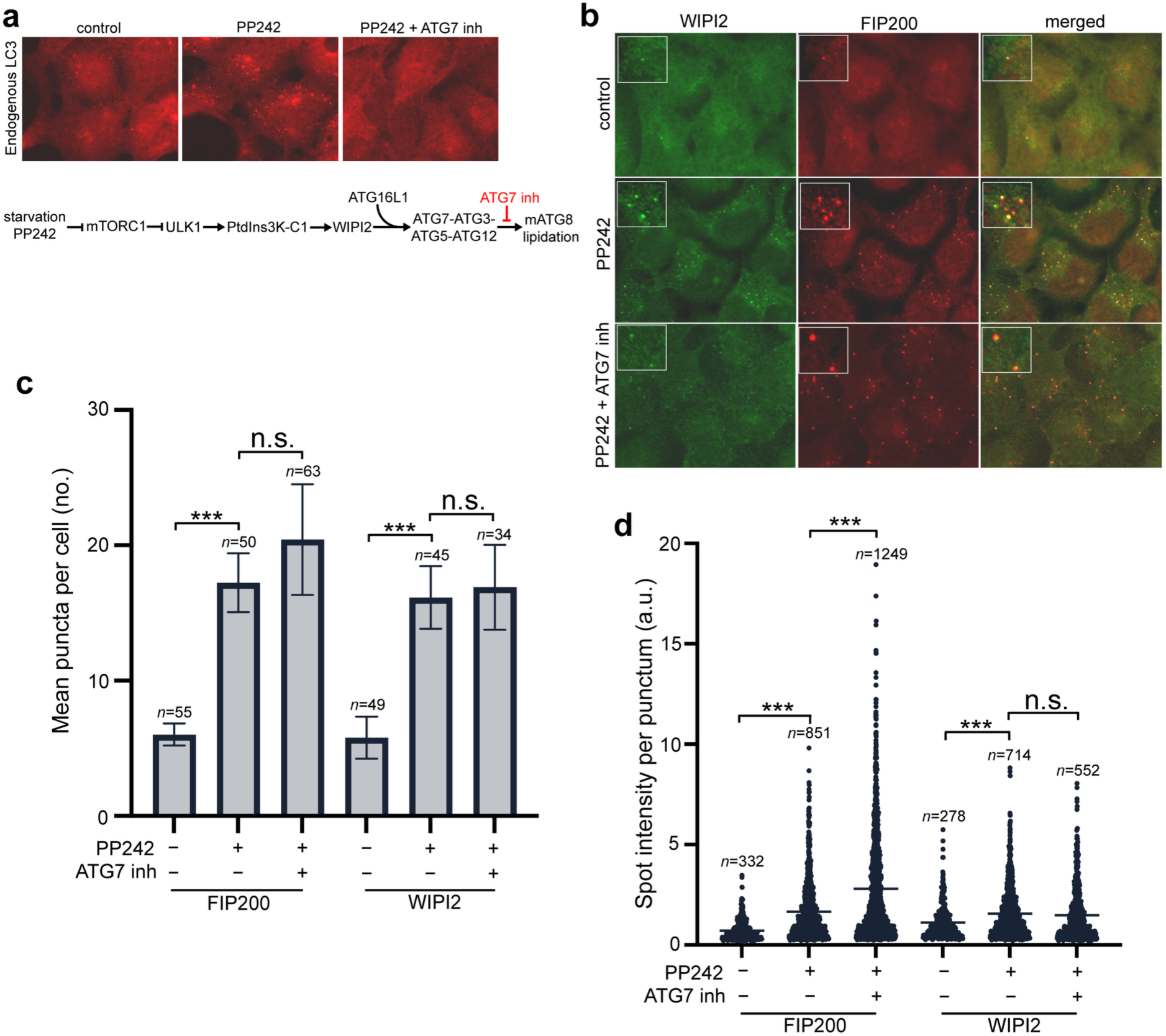
PtdIns3P production is reduced by inhibiting its downstream mATG8 lipidation machinery in cells. **a** HEK293 cells were treated for 60 min with 1 μM of the mTOR inhibitor PP242 in the presence or absence of an ATG7 inhibitor (inh, used as a pretreatment for 30 min) at 5 μM final concentration. The cells were then stained for immunofluorescence with antibodies to LC3. **b** HEK293 cells were treated as in panel a but were double stained with antibodies to endogenous FIP200 and endogenous WIPI2 as shown. Image acquisition and any intensity corrections were done to the same extent for both channels. **c and d** Quantitation of images shown in panel b. Five fields of each treatment were photographed and the number (**c**) and intensity (**d**) of puncta per cell were estimated using ImageJ. Averages for *n* total cells were compared in a two-way ANOVA with Tukey’s correction for multitesting (ns: *p* > 0.05, ***: *p* < 0.0001).

### GABARAP and other lipidated mATG8s bind to and activate PtdIns3K-C1

Three LIRs have been previously characterized on the PIK3C3, ATG14 and BECN1 subunits of PtdIns3K-C1, with a selectivity for the mATG8s GABARAP and GABARAPL1^23^. However, this study used peptides from LIRs of PIK3C3, ATG14, and BECN1 to characterize their interaction with mATG8s, instead of the full PtdIns3K-C1 assembly. To determine whether the proposed binding between PtdIns3K-C1 subunits and mATG8s does happen within the full complex, we immobilized recombinant mCherry−PtdIns3K-C1 on beads and incubated it with mCerulean-fusions of each of the six human recombinant mATG8s (Fig. 2a). Under a confocal microscope we could see clear binding of all mATG8s to PtdIns3K-C1 and confirmed stronger binding of PtdIns3K-C1 to the mATG8s GABARAP and GABARAPL1 (Fig. 2b). Titration gave an apparent Kd of 1.2 µM for the interaction between PtdIns3K-C1 and GABARAP (Fig. 2c, Supplementary Fig. 1a). This binding was not caused by non-specific interactions with mCerulean or mCherry (Supplementary Fig. 1b). Since mATG8s are covalently coupled to lipids on the growing phagophore in cells, we reconstituted this in vitro using recombinant ATG7 (E1), ATG3 (E2), the ATG12-ATG5–ATG16L1 (E3) complex^17^, and all six human mCerulean-mATG8 proteins on giant unilamellar vesicles (GUVs) (Fig. 2d). This successfully immobilized all mCerulean-mATG8s, but not mCerulean, on GUVs only in the presence of ATP and E1-E2−E3 proteins (Fig. 2e left, mCerulean channel, Supplementary Fig. 1c). These vesicles were incubated with PtdIns3K-C1 to measure the efficiency of the PtdIns3P production in an endpoint assay, using AF647-p40^phox^ PX as a product fluorescent reporter. Interestingly, all lipidated mATG8s except LC3B significantly increased the production of PtdIns3P (Fig. 2e right). GABARAP and GABARAPL1 were the most potent activators of PtdIns3K-C1, which correlated with them having the strongest affinity for PtdIns3K-C1 compared to the other mATG8s (Fig. 2b)^23^. Because GABARAP showed the strongest binding and the highest PtdIns3K-C1 activation, we examined its activation mechanism in a time course experiment. To simplify the lipidation system, we immobilized GABARAP on GUVs by incubating a cysteine-introduced GABARAP mutant (L117C) with DOPE-maleimide−containing GUVs (Fig. 2f). We obtained a clear activation of PtdIns3K-C1 by maleimide-lipidated GABARAP, to the same extent as the enzymatic lipidation in endpoint assays (Supplementary Fig. 1d), whereas soluble GABARAP (not membrane-bound) was not able to activate PtdIns3K-C1 (Supplementary Fig. 1e). The time course experiment showed that membrane-anchored GABARAP dramatically increased the initial rate up to 14-fold (Fig. 2g). Taken together, these results suggest that the binding of PtdIns3K-C1 to the lipidated mATG8s results in PtdIns3K-C1 activation on membranes. For the rest of this study, we focused on GABARAP as a model of an mATG8 able to activate PtdIns3K-C1, considering its high affinity and activation potency for the complex among all six mATG8s.

**Figure 2.**
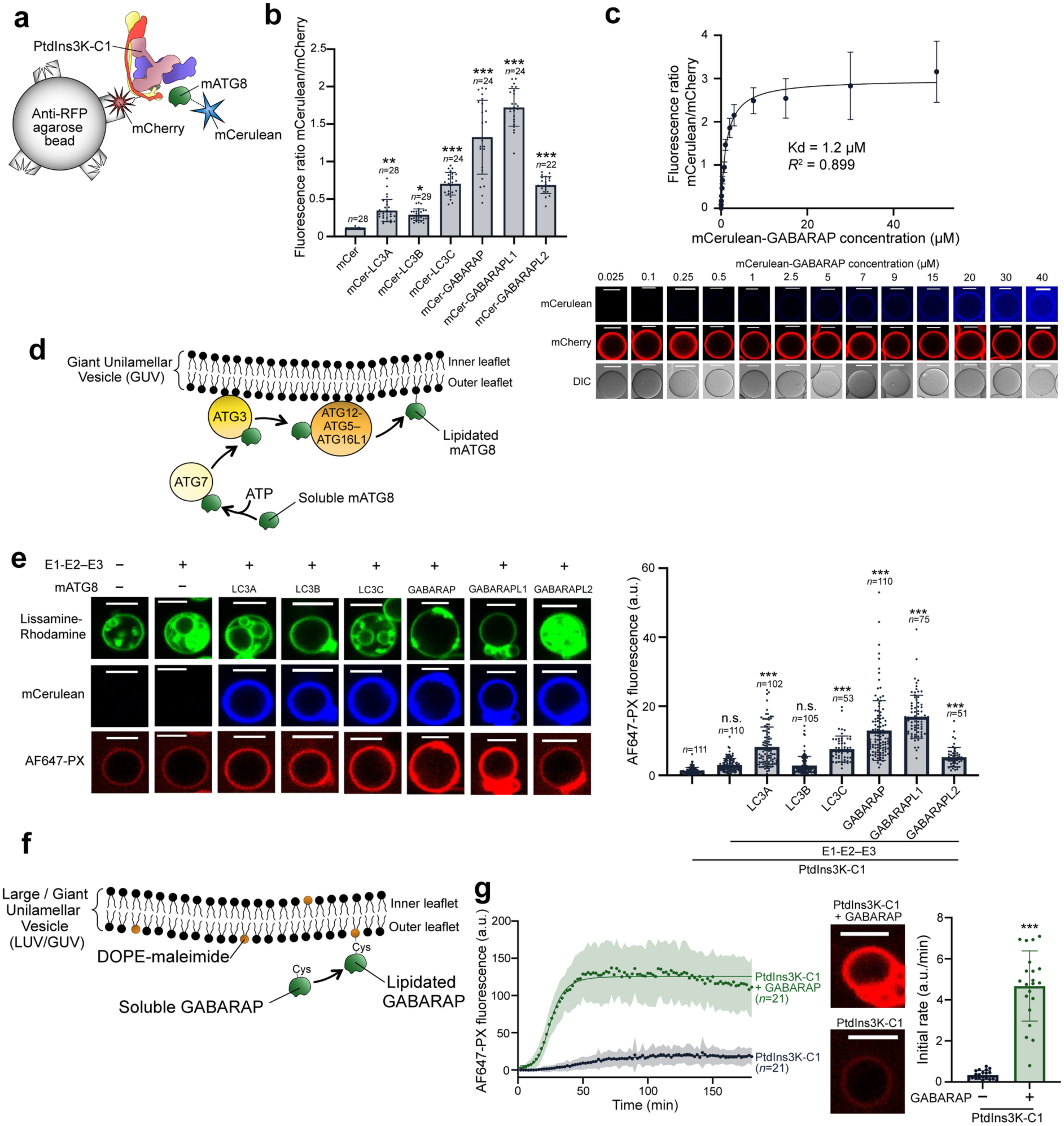
mATG8s bind to and activate PtdIns3K-C1. **a** Design of the beads-based binding assay. RFP-trap agarose beads were coated with mCherry-tagged PtdIns3K-C1, incubated with different mCerulean (mCer)-tagged mATG8 proteins and imaged by confocal microscopy. **b** Quantification of the mCerulean (mCer)-tagged mATG8 proteins binding to PtdIns3K-C1. The plot shows the average for *n* beads ± *SD* and is representative of three independent repeats of the experiment. Means were compared to the mCerulean control with a one-way ANOVA with Tukey’s correction for multitesting (n.s.: *p* > 0.05, *: *p* < 0.05, **: *p* < 0.002, ***: *p* < 0.0001). **c** Quantification of the interaction between PtdIns3K-C1 and different concentrations of mCerulean-GABARAP. Top: The plot shows the average of at least 29 beads ± *SD* and is representative of three independent repeats of the experiment (Supplementary Fig. 1a). Bottom: Representative confocal micrographs of the experiment in the top panel. Scale bars: 50 µm. **d** In vitro reconstitution of the mATG8 enzymatic lipidation. Vesicles were incubated with ATG7 (E1), ATG3 (E2), ATG12-ATG5−ATG16L1 (E3), different mATG8s and ATP. **e** Activation of PtdIns3K-C1 by lipidated mATG8s on GUVs. E1-E2−E3 represent the lipidation enzymes listed above. Left: Representative confocal micrographs of GUVs, showing Lissamine-Rhodamine (lipids), mCerulean (mATG8) and AF647 (PtdIns3P) channels. Scale bars: 5 µm. Right: Average of the AF647-PX fluorescence intensity in arbitrary units (a.u.) for *n* GUVs ± *SD.* Means were compared to the PtdIns3K-C1 only control with a two-way ANOVA with Tukey’s correction for multitesting (n.s.: *p* > 0.05, ***: *p* < 0.0001). **f** Chemical lipidation of GABARAP using a maleimide reaction. Cys = cysteine. **g** GUV-based time course PtdIns3K-C1 activity assay with or without maleimide-lipidated GABARAP, for one representative experiment of 21 repeats. Left: PtdIns3P production over time expressed in average AF647-PX fluorescence ± *SD* for *n* GUVs (in arbitrary units a.u.). Middle: Representative micrographs of GUVs at the end of the reaction in the AF647 channel. Scale bars: 5 µm. Right: Initial rates of the reaction curves in the left panel, expressed in AF647-PX fluorescence intensity/min. The means were significantly different in a two-tailed unpaired *t*-test (***: *p* < 0.0001).

### GABARAP binds PtdIns3K-C1 through an atypical bipartite interaction

To gain insight on which sites are responsible for the GABARAP interaction in the full complex assembly, we solved a 3.6 Å cryo-EM structure of PtdIns3K-C1 with the MIT domain of NRBF2 bound to soluble GABARAP (Fig. 3a). A beads-based interaction assay confirmed that NRBF2 MIT does not affect the binding of GABARAP to PtdIns3K-C1 (Supplementary Fig. 1b). We observed an additional density at the N-termini of the ATG14 and BECN1 subunits that clearly fitted one GABARAP molecule (Fig. 3a and b). Two interaction sub-sites with GABARAP could be distinguished: one on ATG14 (NA), and one on BECN1 (NB), both NA and NB sites forming the GABARAP-binding N site (Fig. 3c). The NB site showed density for the LIR of BECN1 97-FTLI-100 contacting the LDS formed by hydrophobic pockets (HP) HP1 and HP2 in GABARAP and we built a model for the interaction guided by the previously described crystal structure of GABARAP bound to the BECN1 LIR (Fig. 3d)^23^. Whereas the NB site forms a canonical LIR-LDS interaction with GABARAP, the contacts we saw between GABARAP and the NA site have no precedent. In presence of GABARAP, the ATG14 N-terminus showed an additional density compared to the previous PtdIns3K-C1 structure^35^, suggesting its interaction with GABARAP stabilized this flexible region. In the NA site we saw ATG14 residues V38 and A39 making clear hydrophobic contacts with the GABARAP ɑ2-β1 loop residues K24 and P26, respectively. Although LIR peptide/LDS interactions have been well characterized^36^, this GABARAP ɑ2-β1 loop-mediated interaction with an mATG8 is the first ever to be described. Two CXXC zinc fingers could be modelled in the vicinity of the NA site (Fig. 3d), one involving BECN1 C137 and C140 and ATG14 C43 and C46, and another with BECN1 C18 and C21 and ATG14 C55 and C58. Interestingly, these BECN1 cysteines were reported in PtdIns3K-C2 to form an intramolecular zinc finger^37^ but are seen here in PtdIns3K-C1 in two intermolecular zinc fingers. Our results also show that FIP200 is not required for the zinc fingers in the N-termini of BECN1 and ATG14 to be ordered, contrary to a previous claim^38^. For the other two previously reported GABARAP binding sites in PtdIns3K-C1 subunits^23^, the ATG14 LIR 435-WENL-438 was not observed in our density as it belongs to a long flexible region from residues 402 to the C-terminus of ATG14 that cannot be resolved in our structure (see below); and the PIK3C3 LIR 250-FELV-253 was seen inserted into a pocket of PIK3R4 residues including R46, N79, Q88, K89 and E334 that likely hold the LIR and thus prevent any binding to an mATG8 (Fig. 3e). Taken together, our cryo-EM structure revealed that GABARAP binds PtdIns3K-C1 at the N-termini of ATG14 and BECN1. The BECN1 LIR and residues V38 and A39 of the ATG14 N-terminus are respectively engaging the LDS of GABARAP and its ɑ2-β1 loop in a newly discovered bipartite interaction.

**Figure 3.**
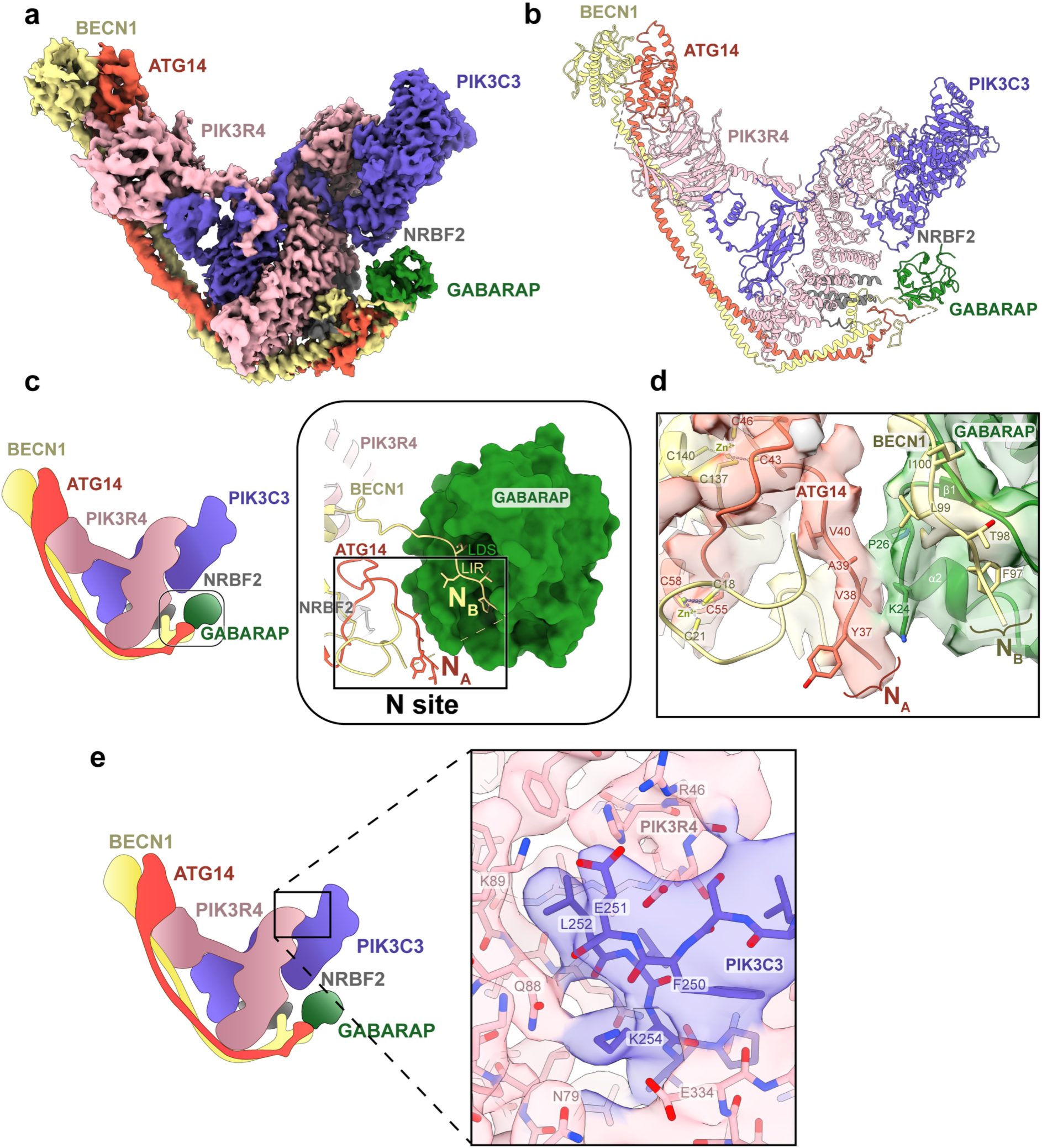
PtdIns3K-C1 binds GABARAP via an atypical bipartite interaction. **a** Cryo-EM composite density map of PtdIns3K-C1−NBRF2 MIT domain bound to GABARAP (main conformation, EMD-56357). A composite map from 5 local refinements was built at 3.58 Å resolution (resolution of the consensus map). **b** Atomic model of PtdIns3K-C1−NRBF2 MIT domain bound to GABARAP (main conformation, PDB 9TW2). **c** Close-up view of the bipartite interaction between PtdIns3K-C1 N site and GABARAP. GABARAP is represented as a surface, and its interactions with ATG14 37-40 and BECN1 97-100 are defined and labelled as NATG14 (NA) and NBECN1 (NB) sites, respectively, both constituting the N site. The NB site consists of the LIR of BECN1, interacting with the LDS of GABARAP. **d** Close-up view of the residues involved in GABARAP binding in NA and NB sites, and neighbouring zinc fingers. The interaction between PtdIns3K-C1 and GABARAP involves ATG14 residues 37 to 40 (NA), BECN1 residues 97 to 100 (NB), and GABARAP residues 24 and 26 on the ɑ2-β1 loop. Helix ɑ2 and β-strand β1 are indicated on the GABARAP model. Two CXXC zinc fingers are seen, one Zn2+ ion being coordinated by BECN1 C18 and C21 and ATG14 C55 and C58, and another Zn2+ by BECN1 C137 and C140 and ATG14 C43 and C46. **e** The PIK3C3 LIR is not accessible to GABARAP due to extensive contacts with PIK3R4. Close-up view of the PIK3C3 250-253 LIR in the cryo-EM structure, showing PIK3C3:PIK3R4 contacts between residues pairs E251:R46, L252:Q88, L252:K89 and K254:Q88, respectively.

Overall, the PtdIns3K-C1 structure adopted a similar conformation to the one previously described for PtdIns3K-C1 bound to RAB1A^35^. However, a focused 3D classification on the adaptor arm of PtdIns3K-C1 identified a minor class of particles (98,630 out of 403,177) that showed a translation of up to 7 Å of the end of the adaptor arm made of the BARA domain of BECN1 and the C-terminal domain of ATG14 (Fig. 4a). We described this translation of the adaptor arm as the ‘alternative’ conformation, in opposition to the ‘main’ conformation of PtdIns3K-C1 adaptor arm. This alternative conformation was not observed in the published RAB1A-bound PtdIns3K-C1 structure^35^ and could be induced by GABARAP to facilitate membrane binding allowing the activation of the complex. We could not see any class of particles resembling the ‘active’ conformation as described in Cook, et al.^35^ (PDB 9MHH), and this was also the case for another cryo-EM dataset we collected with PtdIns3K-C1 in presence of the NRBF2 MIT domain and ADP-MgF3 as an ATP transition-state mimic (Fig. 4b)^39^. However, in this nucleotide-containing dataset we clearly saw a distinct ADP density in the ATP-binding pocket, along with a Mg^2+^ ion coordinating the phosphates of the ADP and PIK3C3 residues N748 and D761, making this structure the first report of a nucleotide-bound PIK3C3. PIK3C3 residues from the ATP-loop (P-loop, S614), hinge (F684), the catalytic loop (D743, R744, H745) and the activation loop (D761) are all visible in this structure, as previously reported for other PI3K-family members^40,41^. To compare models of the kinase domains from datasets in the presence and absence of nucleotide, we aligned them on the PIK3C3 C-lobe (686–887), and this gave a root mean square deviation (RMSD) of 0.77 Å and an angle of 0.68° between the N-lobes (553–682). Remarkably, the conformation of the PIK3C3 kinase is identical in both our datasets with or without a nucleotide, similarly to what was previously seen in DmVps34 with or without inhibitors^42^.

**Figure 4.**
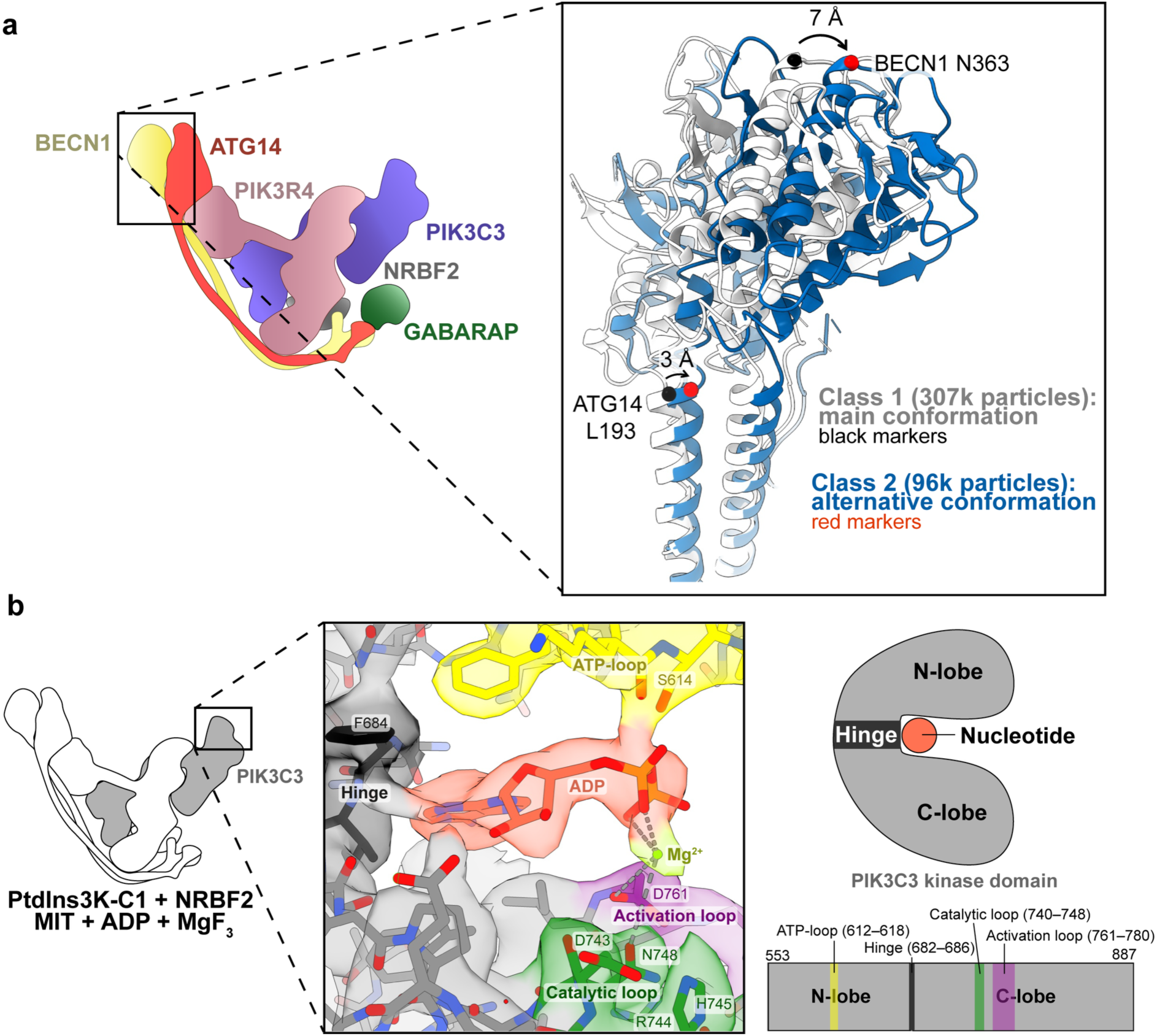
Structural features of the activated PtdIns3K-C1. **a** PtdIns3K-C1 adaptor arm conformation change induced by GABARAP. Two classes were obtained in a 3D focused classification on the adaptor arm of PtdIns3K-C1: the ‘main’ conformation class 1 (white) and the ‘alternative’ conformation class 2 (blue). Black and red markers (respectively) were positioned on the structures to estimate the distances of the shift. **b** Left and middle: Cryo-EM density (EMD-56458) and model (PDB 9TZ3) for PtdIns3K-C1−NRBF2 MIT domain bound to ADP-MgF3, with a close-up view on the nucleotide binding pocket. PIK3C3 ATP-loop (P-loop), hinge, catalytic loop and activation loop are visible around the ADP nucleotide. Mg2+ is coordinated by PIK3C3 residues N748 and D761. Right: PIK3C3 kinase domain organization with the ATP-loop, hinge, catalytic loop and activation loop.

### Identification of another GABARAP binding site with structural mass spectrometry

Because the ATG14 LIR^23^ is part of a flexible region that we did not observe by cryo-EM, we carried out a hydrogen/deuterium exchange coupled to mass spectrometry (HDX-MS) as an orthogonal approach to cryo-EM to investigate the interaction between PtdIns3K-C1 and GABARAP in solution. Upon comparing the exchange profiles between PtdIns3K-C1 in the presence of GABARAP and a control without GABARAP, two main regions in PtdIns3K-C1 showed significant HDX changes upon GABARAP binding (Fig. 5 and Supplementary Fig. 2a). The first one is localized in the C-termini of BECN1 and ATG14 (hereafter called the C site), where strong protections were observed for residues spanning the ATG14 LIR (residues 429−443) and the BECN1 BARA domain (residues 278−285). The second region involved the GABARAP-binding N site of PtdIns3K-C1 that we characterized by cryo-EM, with the NA site showing protection (residues 39−48), while NB LIR site was not covered in our HDX-MS dataset. Additionally, we observed protection in the N-termini of BECN1 (residues 19−31 and 125−139) and ATG14 (residues 51−56), close to the N site. Interestingly these regions and the ATG14 39−48 region are involved in the intermolecular zinc fingers identified in cryo-EM, suggesting the potential involvement of these zinc fingers in the stabilization of the interaction with GABARAP.

**Figure 5.**
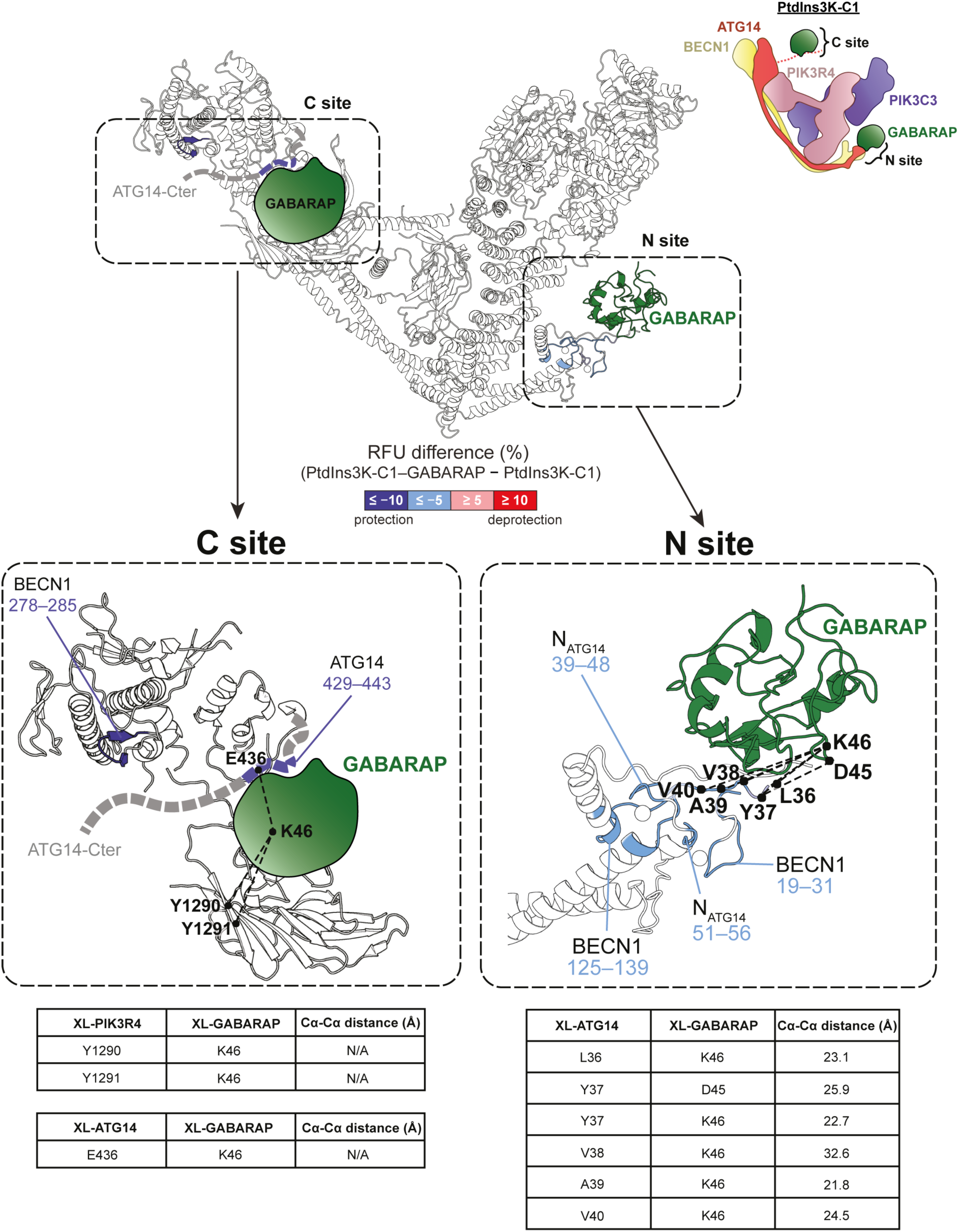
Identification of two GABARAP-binding sites on PtdIns3K-C1 by a combination of structural approaches. HDX changes (relative fractional uptake difference [PtdIns3K-C1−GABARAP − PtdIns3K-C1]) and identified inter-crosslinked sites between GABARAP and PtdIns3K-C1 are shown on PtdIns3K-C1−GABARAP structure (PDB 9TW2). Strong (≤ −10%) and light (≤ −5%) HDX protections are shown in dark and light blue, respectively, and inter-crosslinked sites are represented by dotted lines. Two GABARAP-binding sites were identified in PtdIns3K-C1, the N site with ATG14 and BECN1 N-termini, and the C site spanning the ATG14 LIR motif and BECN1 BARA domain, both sites showing protected regions and inter-crosslinked sites. The two Zn2+ ions of the N site zinc fingers are represented as spheres. HDX-MS difference plots and crosslinking-MS maps are shown in Supplementary Fig. 2.

Next, we used crosslinking coupled to mass spectrometry (XL-MS) to identify spatial proximities between PtdIns3K-C1 and GABARAP, by using succinimidyl-ester diazirine (SDA). Firstly, crosslinks were identified in the C site involving GABARAP residue K46 with ATG14 LIR residue E436 and PIK3R4 residues Y1290 and Y1291 (Fig. 5 and Supplementary Fig. 2b). Interestingly, ATG14 residue E436 is part of the protected region identified in HDX-MS upon GABARAP binding. Both HDX-MS and XL-MS results suggest that PtdIns3K-C1 presents a second binding site involving ATG14 LIR (the C site). Secondly, crosslinks were identified in the N site, involving GABARAP residues D45 and K46 and ATG14 residues 36 to 40 of the NA site, showing protection in our HDX-MS data and directly involved in GABARAP binding in cryo-EM. To ensure that both the N site and the C site can bind GABARAP when it is lipidated, we conducted another crosslinking experiment with GABARAP lipidated on LUVs using maleimide coupling (Fig. 2f, Supplementary Fig. 3). Crosslinks were identified in both the C site and the N site of PtdIns3K-C1, confirming these two regions can bind GABARAP when it is covalently coupled to membranes. Altogether, our data suggest that both the N site (made of NA and NB sites) and the C site (made of the ATG14 LIR) can bind GABARAP, either as a soluble protein or in its lipidated form.

### Mutagenesis of the C site and the N site to disrupt GABARAP binding

To examine the respective roles of the N and C sites on the binding to PtdIns3K-C1, we mutated residues involved in the interaction with GABARAP. The NB and C sites are canonical LIRs, for which the mATG8-binding disrupting mutations are well characterised (the Θ-X-X-Γ LIR motif mutated to A-X-X-A)^23,43^. For the C site, we thus mutated 435-WENL-438 into 435-AENA-438 (hereafter called C^mut^), or we deleted the whole BATS domain carrying this region (ΔC) for the bead-binding assays (see below). For the NB site we mutated 97-FTLI-100 into 97-ATLA-100 (NB^mut^). To disrupt the hydrophobic contacts of GABARAP with the NA site, we deleted the whole ATG14 YVAV motif (residues 37 to 40, ΔNA). We first conducted bead-binding assays using ATG14-BECN1 heterodimers carrying these mutants (Fig. 6a), since all the binding sites we identified are only in these two subunits out of four in PtdIns3K-C1. For each heterodimer, a Kd between mCerulean-GABARAP and mCherry-ATG14−BECN1 was estimated (Fig. 6b, Supplementary Fig. 4a). We only tested the N site mutants in combination with the ΔC mutant as the C site has the strongest affinity for GABARAP and thus masked the effect on the other sites^23^. While it appeared that mutating any site (C, NA or NB) affected the binding to GABARAP, the strongest effects were observed with the ΔC ΔNB^mut^ and ΔC ΔNA NB^mut^ combinations, both increasing the Kd for GABARAP from 1.8 µM for the wild type ATG14-BECN1 heterodimers to about 28 µM that likely corresponds to a non-specific interaction with GABARAP. Taken together, these results confirm that the mutagenesis strategy we applied to the different GABARAP-binding sites was successful and can eliminate binding of PtdIns3K-C1 to GABARAP.

**Figure 6.**
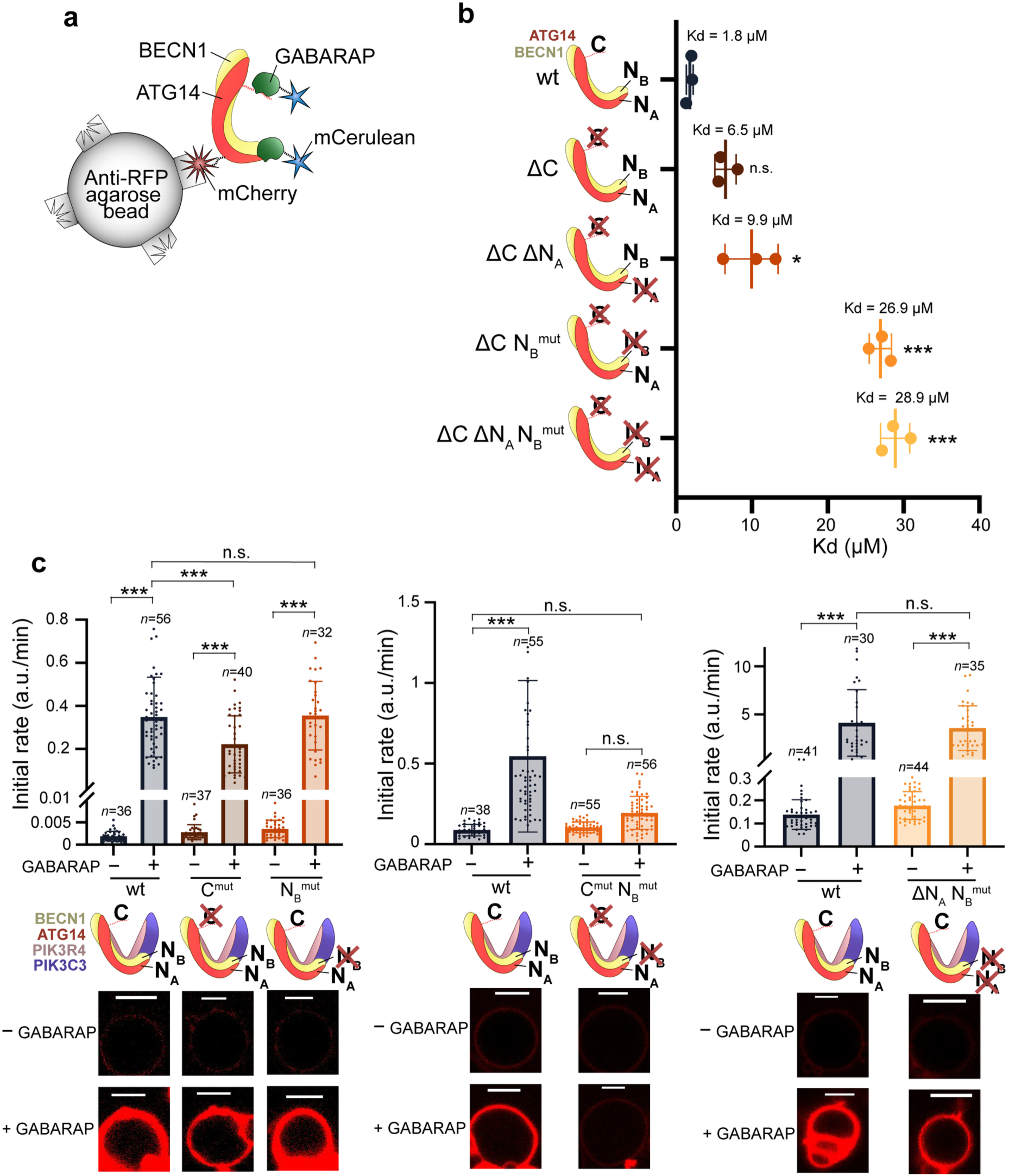
Both C and N sites cooperate for PtdIns3K-C1 activation by GABARAP. **a** Design of the beads-based binding assay. RFP-trap agarose beads were coated with mCherry-ATG14−BECN1 heterodimers, incubated with different concentrations of mCerulean-tagged GABARAP and imaged by confocal microscopy. **b** Affinity for GABARAP of different C site and N site mutants. All titrations are shown in Supplementary Fig. 4a. The plot shows the average ± *SD* of three Kd values determined for repeats of the experiment. Means were compared to the wild type control with a one-way ANOVA with Tukey’s correction for multitesting (n.s.: *p* > 0.05, *: *p* < 0.05, ***: *p* < 0.0001). **c** GUV-based activity assay of full PtdIns3K-C1 assembly carrying C site and N site mutants in presence or absence of maleimide-lipidated GABARAP. Top: PtdIns3P production initial rates expressed in AF647-PX fluorescence/min, average ± *SD* for *n* GUVs. Averages were compared in a two-way ANOVA with Tukey’s correction for multitesting (ns: *p* > 0.05, ***: *p* < 0.0001). Left, middle and right panels correspond to separate experiments, representative of three independent repeats each (Supplementary Fig. 4b). Bottom: Representative micrographs of GUVs at the end of the reaction in the AF647 channel. Scale bars: 5 µm.

### The C and N sites cooperate for PtdIns3K-C1 activation by GABARAP

The mutant combinations in Fig. 6b were incorporated into the full PtdIns3K-C1 assembly and activation assays were done in the presence of lipidated GABARAP on GUVs during time courses (Fig. 6c, Supplementary Fig. 4b). Mutating the ATG14 LIR in the C site (C^mut^) halved the initial rate of GABARAP activation, without abolishing it completely. Mutating the BECN1 LIR (NB^mut^) with or without a deletion of the ATG14 YVAV motif (ΔNA) had no effect on the activation of PtdIns3K-C1 by GABARAP, however a combination of both C and NB mutations completely abolished the ability of GABARAP to activate PtdIns3K-C1. Thus, the NB site mutation had an observable effect on activity only when combined with a mutation of the C site, which is in line with the latter having the greatest affinity for GABARAP (see previous section). In the absence of GABARAP all PtdIns3K-C1 constructs had the same basal PtdIns3P production, confirming none of these mutations affected the integrity of the complex. Taken together, these results functionally validate the interactions sites between PtdIns3K-C1 and GABARAP we identified and indicate that both the C site and the N site in PtdIns3K-C1 are important for GABARAP to fully activate the complex, with the C site being the main contributor to the activation and the resulting positive feedback loop.

## Discussion

In this study, we showed that among the three LIR motifs in PtdIns3K-C1 that were originally proposed to bind GABARAP^23^, only the ones in BECN1 (NB site) and ATG14 (C site) are actually accessible to GABARAP in the full PtdIns3K-C1 assembly, whereas the PIK3C3 LIR is occluded due to numerous interactions with the PIK3R4 protein as seen in our cryo-EM structure (Fig. 3e). However, it is still possible that this region could bind an mATG8 if a conformational change would occur to allow the PIK3C3 LIR to dissociate from PIK3R4, induced, for instance, by the phosphorylation of PIK3C3 S249 by ULK1^23^, or by a change in the nucleotide-binding region of PIK3R4 located near the LIR. An alternative ‘active’ conformation of PtdIns3K-C1 was recently proposed based on a low-resolution class from a cryo-EM study^35^. In this conformation, the PIK3C3 helical domain/kinase unit (FATKIN) rotates by about 140° from the conformation that we observed in our PtdIns3K-C1 structures. Even in this alternative conformation, the interactions with PIK3R4 that occlude the PIK3C3 LIR from binding an mATG8 would persist. In addition to the C and NB LIR-mediated binding sites, our MS and cryo-EM results identified a novel GABARAP binding site NA in the N-terminus of ATG14 forming hydrophobic contacts with the ɑ2-β1 loop of GABARAP (Fig. 3d). Such a bipartite interaction of a protein with an mATG8 is atypical and unique to PtdIns3K-C1. Interestingly, the N-terminal region of the mATG8s is an evolutionarily conserved feature in the mATG8 family that distinguishes these proteins from other ubiquitin-like proteins^44^ and is a region with multiple roles ranging from p62 interaction^45^ to membrane tethering^46,47^.

Our cryo-EM structure of PtdIns3K-C1 with GABARAP provides structural details about the interaction between PtdIns3K-C1 and GABARAP in their soluble form, however, we could not determine the structure of the complex on reconstituted membranes with lipidated GABARAP. Future work will be needed to fully understand how PtdIns3K-C1 is recruited and activated by GABARAP on membranes and to determine whether PtdIns3K-C1 interaction with GABARAP affects GABARAP orientation on membranes^48^. The transition state of a kinase occurs with both ATP and phosphoacceptor bound to the enzyme. There is no structure of PtdIns3K-C1 with either ATP or the PtdIns substrate, despite recent attempts to capture the complex in an activated conformation^35,49^, and a structure of the similar complex PtdIns3K-C2 bound to membranes^15^. Our cryo-EM structures of PtdIns3K-C1 bound to GABARAP or to the nucleotide ADP-MgF3 both have conformations that resemble the ‘inactive’ state described previously for PtdIns3K-C1^35^, however, they provide structural insights into two essential components of PtdIns3K-C1 activation: the binding of a nucleotide and the recruitment of PtdIns3K-C1 to membranes by GABARAP^50^. We could see for the first time at the side chain level how the PIK3C3 kinase active site engages with a nucleotide. In addition, our work paves the way for future efforts aimed at resolving a structure of PtdIns3K-C1 bound to a nucleotide and the lipid substrate on membranes, helped by an activating protein like GABARAP.

We found that the mATG8s GABARAP and GABARAPL1 are more potent activators of PtdIns3K-C1 than members of the LC3 family, in particular LC3A and LC3B. We propose that this selectivity of PtdIns3K-C1 for the GABARAP family of mATG8s arises from the specific residues at the contact of the NA site we discovered. The GABARAP-NA interaction we saw in cryo-EM consists of hydrophobic contacts between residues 24-KYPD-27 of GABARAP and residues 37-YVAV-40 of ATG14. A multi-sequence alignment of all human mATG8s (Supplementary Fig. 5) shows that the KYPD sequence of GABARAP is conserved among all members of the GABARAP family but is not shared with the LC3 family since LC3A/LC3B have a 26-QHPS/T-29 sequence that is likely to prevent such hydrophobic contacts with the PtdIns3K-C1 NA site. Interestingly, LC3C shows in this site a 32-KFPN-35 sequence closely resembling that of GABARAP, which might explain why among the LC3 family LC3C showed a particularly high affinity for and activation of PtdIns3K-C1 (Fig. 2b and e). In addition, the Y25 of GABARAP is known to be involved in a cation-π interaction with GABARAP R28, an interaction that is conserved in the GABARAP family but lost in the LC3 family members LC3A and LC3B, where this pair of residues is replaced by H27 and K30, respectively^44^. LC3C at these positions too (F33 and K36) is more like the GABARAP family. As this cation-π interaction is in the vicinity of the GABARAP residues interacting with NA in our cryo-EM structure, we suggest this conserved cation-π interaction could be critical to limit the flexibility of GABARAPs and LC3C in this site, stabilising the interaction with the PtdIns3K-C1 NA site. We propose that the PtdIns3K-C1 NA site is a likely origin for the selectivity of PtdIns3K-C1 for GABARAPs and LC3C, explaining why these mATG8s have a greater affinity for PtdIns3K-C1, resulting in a higher PtdIns3K-C1 activation. Our results are in line with the preference for the GABARAP family of mATG8s in core autophagy components^21^ and reminiscent of the activation of ULK1 by GABARAP^51^.

Our finding that mATG8s, in particular GABARAP, can efficiently activate PtdIns3K-C1 changes what was the previously accepted order of events in phagophore nucleation and elongation during autophagy. Because the lack of PtdIns3P fails to recruit Atg8/mATG8 proteins both in yeast and mammals during starvation^11,52^, this PtdIns3K-C1−driven PtdIns3P production > recruitment of WIPIs > Atg8/mATG8 lipidation axis is an evolutionally conserved and is an essential pathway for autophagy. Based on this genetic and molecular hierarchy of recruitment, this process was thought so far to be unidirectional. Our results suggest that once lipidated on the growing phagophore, GABARAP can in turn recruit PtdIns3K-C1 and activate it, creating a positive feedback loop allowing more PtdIns3P to be generated and thus more GABARAP to be recruited and lipidated (Fig. 7). This positive feedback loop could be essential to sustain an efficient PtdIns3P production on the elongating phagophore, especially during the engulfment of large cargoes^53^. The GABARAP−PtdIns3K-C1 loop is in line with recent studies suggesting other positive feedback loops between proteins involved in the initial steps of autophagy, such as PtdIns3K-C1 and WIPI2^17^, ULK1 and PtdIns3K-C1^54^, ULK1 and GABARAPs^55^ and ULK1 and WIPI2^56^. This would suggest that during autophagy many positive feedback loops involving PtdIns3K-C1 and other autophagy components happen concomitantly to drastically increase the PtdIns3P production, allowing a rapid growth of the phagophore. This work for the first time revealed the structural details of the GABARAP−PtdIns3K-C1 positive feedback loop and helps the understanding of autophagy at a fundamental and molecular level.

**Figure 7.**
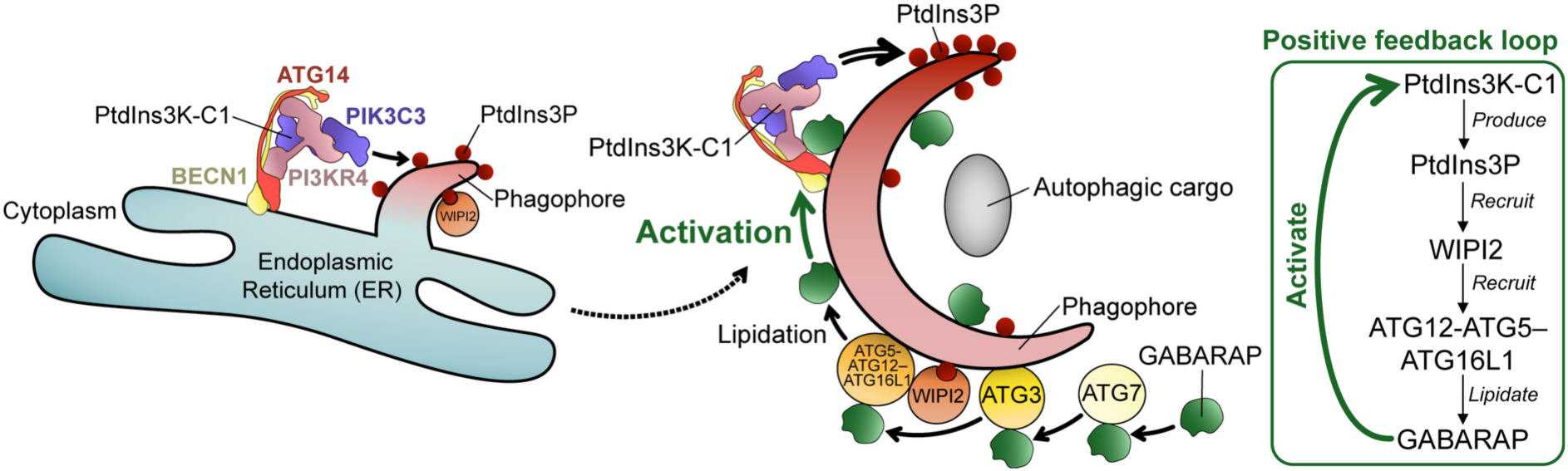
The GABARAP−PtdIns3K-C1 positive feedback loop. During autophagy, PtdIns3K-C1 initially produces PtdIns3P at the ER (left), which can then recruit WIPI2. WIPI2 recruits the ATG5-ATG12–ATG16L1 E3 complex to allow the lipidation of GABARAP on the growing phagophore (middle). GABARAP then recruits PtdIns3K-C1 to the phagophore and activates it, allowing for more PtdIns3P to be produced in a positive feedback loop (right).

## Abbreviations

ATG: autophagy-related
cryo-EM: cryo-electron microscopy
GABARAP: GABA type A receptor-associated protein
HDX-MS: hydrogen/deuterium exchange coupled to mass spectrometry
LC3: (microtubule associated protein 1) light chain 3
LDS: LIR-docking site
LIR: LC3-interacting region
PtdIns3P: phosphatidylinositol-3-phosphate
PIK3C3: phosphatidylinositol 3-kinase catalytic subunit type 3
PIK3R4: phosphoinositide-3-kinase regulatory subunit 4
WIPI: WD repeat domain phosphoinositide-interacting protein;
XL-MS: crosslinking coupled to mass spectrometry.

## Concluding remarks and future perspectives

For the first time we showed that mATG8s, downstream factors of PtdIns3K-C1, can increase its kinase activity, forming a positive feedback loop in cells. Our study reveals that PtdIns3K-C1 binds two mATG8s, one on the C site and one on the unexpected bipartite mATG8 binding site at the N-terminus of PtdIns3K-C1 (the N site). Further investigation will be needed to understand how PtdIns3K-C1 binds GABARAP on membranes. We characterized different mutations in PtdIns3K-C1 that abolish the interaction with GABARAP and its ability to activate PtdIns3K-C1, further investigation will be needed to explore their effects on the phagophore nucleation and elongation in cells.

## Acknowledgements

We thank Saulė Špokaitė for the crYOLO trained model and the GUV generation protocol, Jyothi Nagraj for help with quantitation, Leon Murphy and Oliver Florey for the ATG7 inhibitor, Jerome Boulanger for the modified microscope macros, Tomás Pais de Azevedo, Sarah Lecinski and Nick Barry of the MRC-LMB Light Microscopy Facility for helping with confocal microscopy, the MRC-LMB Mechanical Workshop for custom instruments, Bilal Ahsan, Anna Yeates, Grigory Sharov and Giuseppe Cannone of the MRC-LMB EM Facility for help with cryo-EM data collection, Ivan Clayson, Toby Darling and Jake Grimmett for help with scientific computing, and Olga Perisic and Ksenia Rostislavleva for purifying the NRBF2 MIT protein.

## Author contributions

A.N.D., Y.O., M.B., T.E.M., A.N. and M.M. conducted the research. A.N.D., Y.O., M.B., T.E.M., A.N. and M.M. analysed data. A.N.D, Y.O., M.B., N.T.K. and R.L.W. developed the experimental plan. A.N.D., Y.O., M.B., N.T.K. and R.L.W. wrote the original draft. All authors reviewed and edited the draft.

## Material and Methods

### Protein expression and purification

#### Purification of PtdIns3K-C1 (untagged and mCherry-tagged)

PtdIns3K-C1 constructs were expressed and purified as published before^15^. Briefly, 2 L of Expi293F suspension cells (ThermoFisher A14527) were transfected with different combinations of plasmids containing PtdIns3K-C1 subunits (see Table 1). The cell pellet was lysed in 100 mL of buffer containing 50 mM HEPES pH 8.0, 150 mM NaCl, 0.5 mM tris(2-carboxyethyl)phosphine (TCEP), 1% (v/v) Triton-X-100 (Sigma X100-500), 12% (v/v) glycerol, 2 mM MgCl2, 2 protease inhibitor tablets (Roche 05056489001) and 1 mM phenylmethylsulfonyl fluoride (PMSF), then centrifuged at 14,000 g for 30 min in a Ti45 rotor (Beckman Coulter). The supernatant was passed through a 5 µm syringe filter (Sartorius), then mixed with 2 mL of IgG beads (Cytiva Sepharose 6 Fast flow 17096902) and incubated for 3.5 h at 4°C with 8 rpm rotation. The lysate was then transferred to a gravity flow column and washed with 100 mL of wash buffer containing 50 mM HEPES pH 8.0, 150 mM NaCl, 0.5 mM TCEP, 0.1% Triton-X-100, 12% (v/v) glycerol, 5 mM ATP, 50 mM MgCl2 and 4 µg/mL of RNAse A (Sigma 83834), then with 100 mL of TEV buffer containing 50 mM HEPES pH 8.0, 150 mM NaCl and 0.5 mM TCEP. The protein was cleaved overnight with TEV protease in the TEV buffer at 4°C. The next day, the protein was eluted with a buffer containing 20 mM HEPES pH 8.0, 150 mM NaCl, 0.1% CHAPS (Calbiochem 220201) and 1 mM TCEP, then concentrated to about 500 µL using a 100 kDa cut-off Millipore large concentrator (Millipore, UFC901024) for gel filtration on a Superdex 200 10/30 column (Cytiva 17517501). Proteins were concentrated, aliquoted and frozen in liquid nitrogen and stored at −80 °C.

**Table 1:** Plasmids.

| ID | Construct | Backbone | Marker | Purpose | Reference | Addgene # |
| --- | --- | --- | --- | --- | --- | --- |
| pOP796 | 6xHis-HsNRBF2 MIT (1-89) | pOPTH | amp | Cryo-EM | Ohashi, et al.<sup>57</sup> | 235701 |
| pYO1025 | HsPIK3C3 untagged and HsPIK3R4-3xTEV-ZZ | pCAG | amp | Beads binding assay, GUV assays, cryo-EM, HDX-MS, XL-MS | Ohashi, et al.<sup>57</sup> | 235705 |
| pYO1100 | HsATG14, untagged | pCAG | amp | Beads binding assay, GUV assays | This study | 235707 |
| pYO1006 | HsBECN1, untagged | pCAG | amp | GUV assays | Ohashi, et al.<sup>58</sup> | 235724 |
| pYO1645 | GST-PX domain with Cys at 2 and C85S | pOPTG | amp | GUV assays | Buckles, et al.<sup>78</sup> | 235719 |
| pYO1101 | HsBECN1 and HsATG14, both untagged | pCAG | amp | GUV assays, cryo-EM, HDX-MS, XL-MS | Ohashi, et al.<sup>58</sup> | 235708 |
| Addgene 169079 | 6xHis-TEV-HsATG3 | pETDuet1 | amp | GUV assays, enzymatic lipidation | Fracchiolla, et al.<sup>17</sup> | 169079 |
| Addgene 169076 | 10xHis-TEV-HsATG5-HsATG7-HsATG10-HsATG12-HsATG16L1-TEV-StrepII | pGBdest | kan + gen | GUV assays, enzymatic lipidation | Fracchiolla, et al.<sup>17</sup> | 169076 |
| pYO1562 | mCherry-HsBECN1 | pEGFP-C1 | kan | Beads binding assay | this study | 252672 |
| pYO1601 | 6xHis-TEV-mCerulean-HsLC3A (1-120) | pOPTH | amp | Beads binding assays, GUV assays | this study | 252673 |
| pYO1609 | 6xHis-TEV-mCerulean-HsLC3B (1-120) | pOPTH | amp | Beads binding assay, GUV assays | this study | 252674 |
| pYO1610 | 6xHis-TEV-mCerulean-HsLC3C (1-126) | pOPTH | amp | Beads binding assay, GUV assays | this study | 252675 |
| pYO1611 | 6xHis-TEV-mCerulean-HsGABARAP (1-116) | pOPTH | amp | Beads binding assay, GUV assays | this study | 252676 |
| pYO1612 | 6xHis-TEV-mCerulean-HsGABARAPL1 (1-116) | pOPTH | amp | Beads binding assay, GUV assays | this study | 252677 |
| pYO1613 | 6xHis-TEV-mCerulean-HsGABARAPL2 (1-116) | pOPTH | amp | Beads binding assay, GUV assays | this study | 252678 |
| pYO1594 | mCerulean-TEV-14xHis | pOP | amp | Beads binding assays | this study | 252679 |
| pYO1495 | mCherry-TEV-GST | pOP | amp | Beads binding assays | this study | 252680 |
| pYO1499 | ZZ-3xTEV-mATG7 | pCAG | amp | GUV assays, enzymatic lipidation | this study | 252681 |
| pYO1928 | ZZ-TEV-HsBECN1 | pRS424 | amp | Beads binding assays | this study | 252682 |
| pYO1932 | ZZ-3xTEV-HsATG14-mCherry | pRS426 | amp | Beads binding assays | this study | 252683 |
| pYO1933 | ZZ-3xTEV-HsATG14 (1-434)-mCherry | pRS426 | amp | Beads binding assays | this study | 252684 |
| pYO1943 | ZZ-TEV-HsBECN1 (F97A I100A) | pRS424 | amp | Beads binding assays | this study | 252685 |
| pYO1948 | ZZ-3xTEV-HsATG14 (1-434 ΔYVAV)-mCherry | pRS426 | amp | Beads binding assays | this study | 252686 |
| pYO1797 | HsATG14 (W435A L438A), untagged | pCAG | amp | GUV assays | this study | 252687 |
| pYO1845 | HsBECN1 (F97A I100A) | pCAG | amp | GUV assays | this study | 252688 |
| pYO1982 | HsBECN1 (F97A I100A) and HsATG14 (ΔYVAV), both untagged | pCAG | amp | GUV assays | this study | 252689 |
| ADp1 | GST-TEV-HsGABARAP (1-116) | pOPTG | amp | Enzymatic lipidation, cryo-EM, HDX-MS, XL-MS | this study | 252690 |
| ADp19 | 6xHis-TEV-mCerulean-HsGABARAP (L117C) | pOPTH | amp | GUV assays | this study | 252703 |
| ADp17 | GST-TEV-HsGABARAP (L117C) | pOPTG | amp | Maleimide lipidation, XL-MS | this study | 252702 |

#### Purification of mCherry-tagged ATG14-BECN1 heterodimers

All mCherry-tagged ATG14-BECN1 heterodimers in Fig. 6 and Supplementary Fig. 4 were purified using a *Saccharomyces cerevisiae* protein expression and purification system modified from Rostislavleva, et al.^13^ and Ohashi, et al.^57^ The plasmids were transformed into a yeast strain YOY193 (Matα pep4::HIS3 prb::LEU2 bar1:HISG lys2::GAL1/10-GAL4 can1 ade2 ura3 leu2–3,112 trp1 TRX1-ADE2::ade2Δ^13^). For each heterodimer, yeast colonies were grown at 30°C until OD600= 0.8–1.5 in 100 mL of medium lacking uracil and tryptophan (–URA –TRP) (0.67 % yeast nitrogen base without amino acids, 0.5 % casamino acids, 0.002 % adenine sulfate, 0.002 % tyrosine and 0.002 % leucine) supplemented with 2 % glucose. The whole culture was added to 700 mL of –URA –TRP medium supplemented with 2 % galactose, 2 % glycerol, 3 % lactic acid (Merck L4263), and 10 µM ZnCl2, and grown at 30°C for 26 h. Cells were harvested in 50 mL tubes, and lysed in lysis buffer (50 mM Tris at pH 9.1, 150 mM NaCl, 0.5 mM TCEP, 1% Triton-X, 2 mM MgCl2, 0.5 mM phenylmethylsulfonyl fluoride [PMSF], and a protease inhibitor tablet [Roche 05056489001]) with 3 g of 0.5 mm zirconia beads (YTZ® GRINDING MEDIA, Tosh) using a cell disruptor (FastPrep-24, MP Biomedicals) at intensity 6.5 for 40 s. Lysates were spun at 3000 g for 1 min. Supernatants were further spun at 14,000 g for 30 min in a Ti45 rotor (Beckman Coulter). The supernatant was passed through a 5 µm syringe filter (Satorius), then incubated with 0.5 mL of IgG beads (GE Healthcare 17-0969-02) for 3.5 h at 4°C with 8 rpm rotation. The beads/lysate mixture was transferred to a gravity flow column, and washed with 100 mL of wash buffer (50 mM HEPES, pH 8.0, 150 mM NaCl, 0.1% Triton X-100, 0.5 mM TCEP, 5 mM ATP [Sigma, A2383], 50 mM MgCl2, 5 µg/mL RNaseA [Sigma, 83834]) and 100 mL of TEV buffer (50 mM HEPES, pH 8.0, 150 mM NaCl, 0.5 mM TCEP). Five mL of TEV buffer and 10 µL of 4.7 mg/mL TEV protease were added and incubated at 4 °C overnight without rotation. The eluate was collected and the proteins were eluted three more times with 5 mL buffer each. The elution fractions were combined and passed through a 0.45 µm syringe filter (Thermo Fisher 15191499), then concentrated using a 50 kDa concentrator (Millipore, UFC905024) before running on a Superdex 200 10/30 column equilibrated with 20 mM HEPES, pH 8.0, 150 mM NaCl, 0.5 mM TCEP. The main peak fractions were combined and concentrated using a 50 kDa concentrator (Millipore, UFC905024). The protein was aliquoted, frozen in liquid nitrogen, and stored at −80 °C.

#### Purification of mCerulean-mATG8s, mCerulean, and ATG3 proteins

All mCerulean-mATG8 proteins constructs (see Table 1) were designed by truncating the mATG8 C-terminus to expose the last glycine, to allow the enzymatic lipidation. For maleimide reaction (ADp19), the last C-terminal residue of GABARAP L117 was replaced by a cysteine. For all constructs, the plasmids were expressed in E. coli C41 (DE3) bacteria, induced with 0.3 mM IPTG at 16°C for 18−20 hours, then the bacterial pellet was lysed using a sonicator (Sonics VCX-750-110) for 5 min on ice (2 s on, 3 s off, and 60% amplitude) in 50 mL buffer containing 50 mM HEPES pH 8.0, 150 mM NaCl, 12% (v/v) glycerol, 20 mM imidazole, 5 mM 2-mercaptoethanol, a protease inhibitor tablet (Roche 05056489001) and 1 mM PMSF. The lysate was centrifuged at 14,000 g for 30 min in a Ti45 rotor (Beckman Coulter). The supernatant was passed through a 5 µm syringe filter (Sartorius), then incubated with Ni-NTA agarose beads (Qiagen 30210) for 1 hour at 4°C with 8 rpm rotation and was transferred to a gravity flow column. The beads were washed with buffer containing 50 mM Tris pH 8.0, 300 mM NaCl, 12% (v/v) glycerol, 20 mM imidazole and 5 mM 2-mercaptoethanol, and eluted with a buffer containing 50 mM Tris pH 8.5, 20 mM NaCl, 300 mM imidazole, 5 mM 2-mercaptoethanol and 1 mM PMSF. The protein was further purified on a Q column (Cytiva 17115401) in buffer 50 mM Tris pH 9.0, 5 mM 2-mercaptoethanol and eluted with buffer 50 mM Tris pH 9.0, 1 M NaCl and 5 mM 2-mercaptoethanol. The peak fractions were combined and concentrated down to ∼ 500 µL using a 10 k cut-off Millipore large concentrator (Millipore, UFC901024), then the protein was subjected to a gel filtration on a Superdex 75 16/60 column (Cytiva 17-1068-01) in a buffer containing 20 mM HEPES pH 8.0, 150 mM NaCl and 0.5 mM TCEP. Proteins were concentrated, aliquoted and frozen in liquid nitrogen and stored at −80 °C. For the ADp19 construct, the final buffer contained 5 mM TCEP instead of 0.5 mM. ATG3 protein was purified essentially in the same way as above except the plasmid was Addgene 169079 and the Q elution buffer was 50 mM Tris pH 9.0, 2 M NaCl and 5 mM 2-mercaptoethanol.

#### Purification of mCherry

The plasmid pYO1495 was transformed into C41(DE3)RIPL cells. The bacteria were induced in 900 mL of 2TY medium with 0.3 mM IPTG at 20°C for 20 hours. The bacterial pellet was lysed using a sonicator for 5 min on ice (2 s on, 3 s off, and 60% amplitude) in 20 mL of lysis buffer containing 50 mM Tris pH 8.0, 300 mM NaCl, 1 mM DTT, a protease inhibitor tablet (Roche 05056489001) and 1 mM PMSF. The lysate was centrifuged at 14,000 g for 15 min in a Ti45 rotor (Beckman Coulter). The supernatant was incubated with 1 mL of Glutathione Sepharose 4B beads (Cytiva, 17075605) at 8 rpm in the cold room for 1 h. The lysate/beads mixture was transferred to a gravity flow column and washed with 250 mL of wash buffer (50 mM Tris pH 8.0, 300 mM NaCl, 1 mM DTT, and 12% glycerol), and 200 mL of TEV buffer (50 mM Tris pH 8.0, 300 mM NaCl, 1 mM DTT), then 10 mL of TEV buffer with 80 µL of 4.7 mg/mL His6-LIPOYL-TEVS219V was added to the beads and incubated at 4°C overnight without rotation. The eluate was collected and the proteins were eluted three more times with 5 mL tev buffer each. The elution fractions were combined, and imidazole was added to a final concentration of 20 mM, then passed through a 5 mL HisTrap HP column (Cytiva 17524801). The flow-through fraction was concentrated using a 10 kDa cut-off concentrator (Millipore 901024). The concentrated protein was subjected to gel filtration on an S75 16/60 column equilibrated with GF buffer (20 mM Tris 8.0, 150 mM NaCl, 0.5 mM TCEP). The peak fractions were combined and concentrated down to 18 mg/mL (641 µM). The protein was aliquoted and frozen in liquid nitrogen and stored at −80 °C.

#### Purification of GABARAP (1−116) and GABARAP L117C

As for the other mATG8 expressed, the C-terminal glycine of GABARAP was exposed by a ΔL117 deletion (ADp1 in Table 1). For the maleimide reaction, a C-terminal cysteine was added (ADp17). For both constructs, the plasmids were expressed in bacteria (E. coli C41 (DE3)) induced in 900 mL of 2TY medium with 0.3 mM IPTG at 18°C for 18 h. The bacterial pellet was lysed using a sonicator (Sonics VCX-750-110) on ice for 3 min (2 s on, 2 s off, and 60% amplitude) in a buffer containing 50 mM HEPES pH 7.4, 150 mM NaCl, 10 mM Dithiothreitol (DTT), 12% (v/v) glycerol, a protease inhibitor tablet (Roche 05056489001) and 1 mM PMSF. The lysate was centrifuged at 14,000 g for 15 min in a Ti45 rotor (Beckman Coulter) and incubated with Glutathione Sepharose 4B beads (GE Healthcare 17-0756-05) for 1 h at 4°C with 8 rpm rotation, then the beads were washed in a gravity flow column with a wash buffer containing 50 mM HEPES pH 7.4, 300 mM NaCl, 12% (v/v) glycerol and 10 mM DTT. The protein was cleaved with TEV protease overnight to remove the N-terminal GST tag, in a buffer containing 50 mM HEPES 7.4, 300 mM NaCl and 5 mM 2-mercaptoethanol. The TEV protease was removed using Ni-NTA agarose beads (Qiagen 30210), the GABARAP proteins concentrated using a 10 kDa cut-off concentrator (Millipore 901024) and further purified with a Superdex 75 16/60 column (Cytiva 17-1068-01) equilibrated in a buffer containing 20 mM HEPES pH 7.4, 300 mM NaCl and 5 mM DTT. For GABARAP L117C (ADp17), no DTT was present in the gel filtration buffer. The protein was concentrated, aliquoted and frozen in liquid nitrogen and stored at −80 °C.

#### Purification of mATG7 protein

The plasmid pYO1499 at 1.1 mg/L was transfected into 1 L of Expi293f cells in Expi293 Expression Medium (ThermoFisher A1435102) with polyethylenimine (PEI) ‘MAX’ (Polysciences 24765, 1 mg/mL in PBS) at 3 mg/L culture, and grown at 37 °C, 8% CO2, and 125 rpm shaking for 2 days. Cell pellets were suspended in 30 mL lysis buffer (50 mM HEPES, pH 8.0, 150 mM NaCl, 1% Triton X-100 [Sigma, X100], 12% glycerol, 0.5 mM tris[2-carboxyethyl]phosphine [TCEP, Soltec Ventures, M115], 2 mM MgCl2, 1× EDTA-free inhibitor tablet [Roche, 05056489001], and 1 mM PMSF), then spun at 14,000 g for 30 min in a Ti45 rotor (Beckman Coulter). The supernatant was passed through a 5 µm syringe filter (Satorius), then incubated with 2 mL of IgG beads (GE Healthcare 17-0969-02) for 3.5 h at 4 °C with 8 rpm rotation. The beads/lysate mixture was transferred to a gravity flow column, and washed with 150 mL of wash buffer (50 mM HEPES, pH 8.0, 150 mM NaCl, 0.1% Triton X-100, 0.5 mM TCEP, 5 mM ATP [Sigma, A2383], 50 mM MgCl2, 5 µg/mL RNaseA [Sigma, 83834]) and 150 mL of TEV buffer (50 mM HEPES, pH 8.0, 150 mM NaCl, 0.5 mM TCEP). Ten mL of TEV buffer and 30 µL of 4.7 mg/mL TEV protease were added and incubated at 4 °C overnight without rotation. The eluate was collected and the proteins were eluted three more times with 5 mL buffer each. The elution fractions were combined passed through a 0.45 µm syringe filter (Thermo Fisher 15191499), then concentrated using a 50 kDa concentrator (Millipore, UFC905024) before running on an Superose6 10/30 column equilibrated with 20 mM HEPES, pH 8.0, 300 mM NaCl, 0.5 mM TCEP. The main peak fractions were pooled and TCEP was added to a final concentration of 2 mM, then concentrated using a 50 kDa concentrator at 17.57 mg/mL (225 µM). The protein was aliquoted and frozen in liquid nitrogen and stored at −80 °C.

#### Purification of ATG5-ATG12–ATG16L1 complex

The Addgene 169076 plasmid was transformed into DH10EMBacY cells, and white colonies were selected on LB agar containing kanamycin, gentamycin, tetracycline, Bluo-gal, and IPTG for bacmid purification. The generated bacmid was transfected into Sf9 cells at a cell density of 1.2 x10^6^ using FuGENE HD (Promega No E2311) and grown in 6-well plates in Insect XPRESS w/ L-Gln (Lonza LZBE12-730Q) at 27°C for 6 days for virus production. The virus was used to infect 1 L of Sf9 cells. Cells were grown for 2 days at 27°C. The cell pellets were suspended in 75 mL of lysis buffer (25 mM HEPES pH 7.4, 300 mM NaCl, 1 mM DTT, 2 mM MgCl2, and 12% glycerol, 1× EDTA-free inhibitor tablet (Roche, 05056489001), and 1 mM PMSF), lysed by sonication with 60% power, 2 s on 8 s off for 2 min 30 s. then spun at 14,000 g for 30 min in a Ti45 rotor (Beckman Coulter). The supernatant was passed through a 5 mm syringe filter (Satorius), then passed through a 5 µL strep column (StrepTrap HP, Cytiva 10298754). The column was washed with 50 mL lysis buffer and 100 mL wash buffer (25 mM HEPES pH 7.4, 300 mM NaCl, 1 mM DTT, 2 mM MgCl2, and 12% glycerol), then eluted with 50 mL of elution buffer (25 mM HEPES pH 7.4, 150 mM NaCl, 1 mM DTT, and 10 mM desthiobiotin). The elute was concentrated using a 100 kDa cut-off concentrator (Millipore, UFC901024) for gel filtration using a Superose6 10/30 column equilibrated with gel filtration buffer (25 mM HEPES 7.4, 300 mM NaCl, and 0.5 mM TCEP). The peak fractions were concentrated using a 100 kDa cut-off concentrator (Millipore, UFC901024) at 12.6 mg/mL (108 µM). The protein was aliquoted and frozen in liquid nitrogen and stored at −80 °C.

#### Purification and labelling of p40^phox^ PX domain

Bacteria (E. Coli C41 [DE3]) were harvested after expressing the plasmid pYO1645 upon induction in 900 mL of 2TY medium with 0.3 mM IPTG at 18°C for 19 h. The pellet was sonicated on ice for 5 min (2 s on, 3 s off, and 60% amplitude) in lysis buffer containing 20 mM HEPES pH 8.0, 200 mM NaCl, 1 mM TCEP, 0.05 μL/mL universal nuclease (ThermoFisher 88702) and 0.5 mg/mL lysozyme (MP Biomedicals 195303). The lysate was centrifuged at 14,000 g for 15 min in a Ti45 rotor (Beckman Coulter) and incubated with 2 mL of affinity Glutathione Sepharose resin (GE Healthcare 17-0756-05), washed with 100 mL of wash buffer containing 20 mM HEPES pH 8.0, 300 mM NaCl and 1 mM TCEP) and 100 mL of TEV buffer containing 20 mM HEPES pH 8.0, 200 mM NaCl and 1 mM TCEP. The N-terminal GST tag was cleaved with TEV protease by overnight incubation. Fractions were eluted, concentrated using a 10 kDa Amicon Ultra15 concentrator (Millipore, UFC901024), and loaded on a Superdex 75 16/60 column (Cytiva 17-1068-01) for further purification by gel filtration, in a buffer containing 20 mM HEPES pH 8.0, 200 mM KCl and 1 mM TCEP. The purified PX domain was labelled using the Alexa Fluor 647 (AF647) C2 Maleimide kit (Invitrogen A20347), using a Heparin column (Cytiva 17040701) to separate the labelled protein from the free dye. The protein was concentrated, aliquoted and frozen in liquid nitrogen and stored at −80 °C.

#### Purification of NRBF2 MIT domain

The plasmid pOP796^57^ was expressed in E.coli C41(DE3) cells in 2xTY medium supplemented with 0.1 mg/mL ampicillin, and induced with 0.3 mM IPTG for 12 hours at 20 °C. Bacteria pellets were centrifugated at 6,700 g for 20 min and resuspended in 75 mL lysis buffer (25 mM TRIS pH 7.5, 300 mM NaCl, 10 mM imidazole, 2 mM 2-mercaptoethanol, 0.5 mg/mL lysozyme, 0.05 µL/mL universal nuclease) and sonicated (Sonics VCX-750-110) for 6 min (10 s on, 10 s off, 60% amplitude). The lysate was centrifugated at 142,000 g for 30 min at 4 °C, and the supernatant filtered through a 0.45 µm syringe filter. The sample was loaded onto a 5 mL HisTrap FF column (Cytiva 17-5255-01) equilibrated with wash buffer (20 mM TRIS pH 7.5, 150 mM NaCl, 10 mM imidazole, 1 mM TCEP), washed with 100 mL wash buffer, 50 mL Ni-A buffer (20 mM TRIS pH 7.5, 150 mM NaCl, 10 mM imidazole, 1 mM TCEP), and eluted with a gradient of Ni-B buffer (20 mM TRIS pH 7.5, 150 mM NaCl, 300 mM imidazole, 1 mM TCEP). Peak fractions were pooled, diluted two-fold with Q-A buffer (20 mM TRIS pH 7.5, 0.5 mM TCEP), and applied to a 5 mL HiTrap Q HP column (Cytiva 17115401), washed with 50 mL Q-A buffer, and eluted with a gradient of Q-B buffer (20 mM TRIS pH 7.5, 1 M NaCl, 0.5 mM TCEP). Peak fractions were pooled and concentrated using a 3 kDa concentrator (Millipore, UFC900324) and loaded onto an S75 16/60 gel filtration column (Cytiva 17-1068-01) equilibrated in 20 mM HEPES pH 7.0, 100 mM NaCl, 0.5 mM TCEP. Peak fractions were pooled and concentrated to 136 µM, then aliquoted and frozen in liquid nitrogen and stored at −80 °C.

### Confocal microscopy beads-based binding assays

Essentially, 1 µL of anti-RFP agarose beads (Chromotek RTA-10) was used per well of a 384-well plate (Greiner 781856), and the bead volume was calculated depending on the number of wells. The beads were resuspended three times with 1 mL of buffer containing 25 mM HEPES pH 8.0, 100 mM NaCl, 0.5 mM TCEP pH 7.4 and 0.1 mg/mL BSA (Sigma A7030) and centrifugated at 600 g. Excess buffer was removed, and the beads were incubated with 1.7 µM of mCherry, mCherry−PtdIns3K-C1 or different mCherry-ATG14−BECN1 constructs for 30 min on ice. Some beads were also added 5 µM NRBF2 MIT. Beads were washed 3 times as above, and 3 µL of beads were added to 27 µL of 10 µM mCerulean-GABARAP (1−116) in a 384-well glass bottom plate (Greiner 781856). Reactions were incubated for 30 min on ice in the sealed plate covered from light. Beads were then imaged on an inverted confocal microscope (LSM 780, Zeiss) with a 10x objective (EC Plan-Neofluar 10x/0.3 M27, Zeiss). Beads were visualized using the DIC channel, the mCerulean channel was excited using a 458 nm Argon laser and collected with a band of 463−556 nm, and the mCherry channel was excited using a diode-pumped solid-state (DPSS) 561 nm laser and collected with a 578−696 nm band. Images were taken using ZEN software (2.1 SP3, Zeiss) and analysed with FIJI software (ImageJ2 2.14.0). The surface of beads was selected as an elliptical ROI (regions of interest), and the fluorescence intensity was determined for each channel using a previously described macro^37,58^. Ten background ROI without beads were selected and their corresponding intensity average subtracted to the fluorescence of the beads for each channel. The mCerulean fluorescence intensity was normalized by the mCherry fluorescence for each bead, to account for coating discrepancies on the beads as well as out-of-focus beads. Data was analysed in Prism10 (GraphPad), and binding affinities and corresponding *R*^2^ were determined with the non-linear regression fitting tool with the ‘one site-binding’ option. The negative control with mCherry immobilized on beads showed no binding to mCerulean-GABARAP, nor did mCherrry–PtdIns3K-C1 to mCerulean (Supplementary Fig. 1b).

### GUV preparation

A thin PVA gel was created by depositing 88 µL of 5% w/v polyvinyl alcohol (PVA, Sigma 8.14894, MW approx. 145 kDa) on a 13 mm coverslip (VWR 631-0150), followed by a 30 s centrifugation at 6000 rpm on a microfuge (IKA mini G) using a custom-made coverslip adaptor. The PVA gel was dried at 60°C for 1 h. The lipid stocks of Table 2 were mixed to prepare the lipid mixtures ‘GUV DO base’ at 1 mg/mL in chloroform, as shown in Table 3. The lipid mixture was carefully deposited on the PVA gel by two applications of 7.5 µL, and the remaining chloroform was evaporated by 1 h desiccation under vacuum. The GUVs were swollen by adding 220 µL of swelling solution (500 mM D-Glucose, Fisher 50-99-07) on the coverslips in a 24-well plate (Costar 3526, Corning) and let incubated for 1 h at room temperature. GUVs were then pipetted into a 2 mL tube (previously coated with 5 mg/mL BSA [Sigma A7030] and washed with swelling solution).

**Table 2:** Lipids.

| Lipid | Manufacturer | Catalogue number |
| --- | --- | --- |
| Liver PI (mixed chains, bovine) | Avanti Polar Lipids, Inc. | 840042C |
| DOPS | Avanti Polar Lipids, Inc. | 840035C |
| DOPE | Avanti Polar Lipids, Inc. | 850725C |
| DOPC | Avanti Polar Lipids, Inc. | 850375C |
| Lissamine-Rhodamine-DOPE | Avanti Polar Lipids, Inc. | 810150C |
| DSPE-PEG(2000)-Biotin | Avanti Polar Lipids, Inc. | 880129C |
| DOPE-MCC | Avanti Polar Lipids, Inc. | 780201C |

**Table 3:**
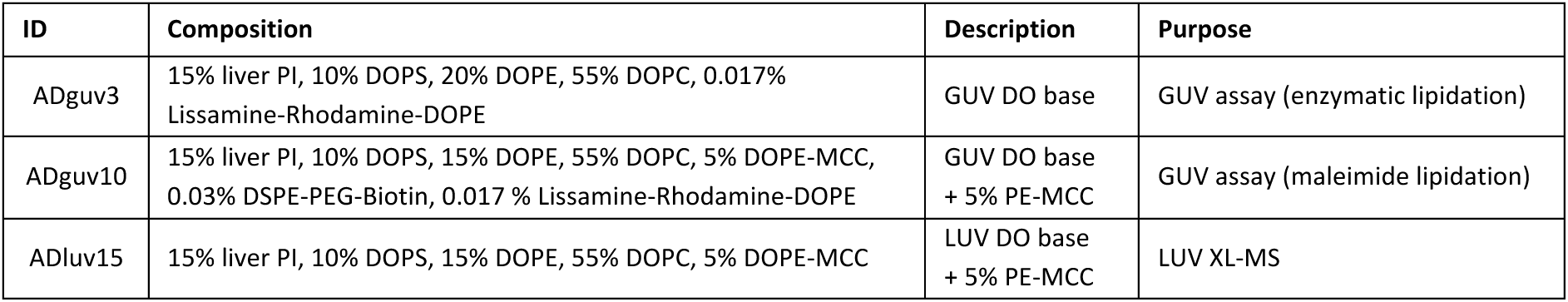
Lipid Mixtures.

### PtdIns3K-C1 activation assay on GUVs

For the GUV enzymatic lipidation, ADguv3 GUVs were added to the wells of a 384-well glass-bottom plate (Greiner 781856), 12 µL GUVs for 50 µL final volume in observation buffer containing 50 mM HEPES pH 7.4 and 271 mM NaCl. ATP-containing buffer was also added, for a final concentration of 25 mM HEPES pH 7.4, 2 mM MnCl2, 1 mM MgCl2, 1 mM TCEP pH 7.4, 1 mM ATP. Proteins were diluted in buffer containing 25 mM HEPES pH 7.4, 150 mM NaCl, 1 mM TCEP pH 7.4 and 0.5 mg/mL BSA (Sigma A7030), and added to the GUV mixture for a final concentration of 20 nM PtdIns3K-C1, 3.5 µM AF647-PX, 200 nM ATG7, ATG3 and ATG12-ATG5–ATG16L1, and 2.5 mM mCerulean-mATG8. The reaction was incubated for 2 h at room temperature.

For the endpoint GUV maleimide lipidation, a 384-well glass-bottom plate (Greiner 781856) was coated with 0.1 mg/mL avidin (Invitrogen A2667) in 1 mg/mL BSA (Sigma A7030)-containing PBS. Wells were washed twice with observation buffer containing 50 mM HEPES pH 7 and 271 mM NaCl, the buffer was removed and 5 µL of 0.1 mg/mL BSA-biotin (Sigma A8549) was added to the wells. Next 10 µM of mCerulean-GABARAP (L117C) and 12 µL ADguv10 GUVs in buffer containing 25 mM HEPES pH 7.0, 150 mM NaCl and 0.5 mg/mL BSA (Sigma A7030) were added, for a total volume of 29 µL. After an overnight incubation at 4°C, the maleimide reaction was quenched with 4 washes of 360 µL of a solution containing 31.8 mM HEPES pH 8.0, 173 mM NaCl, 181.8 mM D-Glucose (Fisher 50-99-07) and 5 mM 2-mercaptoethanol (Aldrich M6250). PtdIns3K-C1 and AF647-PX in the ATP-containing buffer were then added at the same concentrations as described above for a final volume of 50 µL and incubated for 2 h at room temperature for the endpoint assays.

GUVs were imaged on an inverted confocal microscope (LSM 880, Zeiss), using a x63 oil immersion objective (Plan-Apochromat 63x/1.4 Oil DIC M27, Zeiss) and ZEN software (Zeiss). The mCerulean channel was excited with a 458 nm Argon laser and collected with a band of 463−556 nm, the Lissamine-Rhodamine channel was excited with a Diode-pumped solid-state (DPSS) 561 nm laser and collected with a 566−629 nm band, and the AF647 channel was excited with a HeNe 633 nm laser and collected with a 638−755 nm band. Micrographs were analysed in FIJI, where the surface of GUVs was selected as elliptical ROIs (regions of interest) and the fluorescence intensity was determined for each channel using the previously described macro^37,58^. Ten background ROI without GUVs were selected and their corresponding intensity average subtracted to the fluorescence of the GUVs for each channel. Data was analysed in Prism10 (GraphPad) for statistics.

For the time course assays, GUVs were prepared following the GUV maleimide lipidation protocol above, but an 8-well glass bottom chamber (Ibidi 80827) was used instead, for a final volume of 200 µL. A chamber system insert plate (PeCon 000 470) was added to the universal mounting frame to immobilize the 8-well chamber. GUV positions were recorded on ZEN software before adding to the GUVs the PtdIns3K-C1 and AF647-PX proteins in the ATP-containing buffer. Imaging of the GUVs was done as detailed above, except that all the fluorescence channels were tracked for all positions every 2 min over a time course of 180 min total, using the Positions and Time Series options in ZEN software, and Definite Focus.2 (Zeiss) was used to keep the GUVs in focus every 8 positions and every time point. Images were analysed as above, but using the version of the macro designed for time courses^37^. Curves were manually aligned by their linear phases in Microsoft Excel, and the initial slopes calculated with the LINEST function. Data was further analysed in Prism10 (GraphPad) for statistics.

### Cryo-electron microscopy sample preparation and screening

For the structure of PtdIns3K-C1 and GABARAP, PtdIns3K-C1 at a final concentration of 3.6 µM was incubated on ice for 30 min with 3.6 µM NRBF2 MIT domain and 40 µM GABARAP (1−116), in a buffer containing 25 mM HEPES pH 8.0, 100 mM NaCl, 1 mM TCEP pH 7.4, 4 mM CHAPSO (Sigma C3649 lot SLBG1674V) and 0.005% (v/v) Nonidet P-40 substitute (Santa Cruz Biotechnology, sc29102 lot G1015). A sample volume of 3.5 µL was applied on a UltrAuFoil R 1.2/1.3 Au300 grid previously glow-discharged (40 mA, 60 s, air, Edwards Sputter Coater S150B), and plunge-frozen in liquid ethane using a Vitrobot Mk 2 (Thermo Fisher) plunger at 14°C, 100% humidity, with 20 s waiting time, 4 s blotting time, and blot force set to +8. Grids were screened on a Glacios (Thermo Fisher) microscope using a Falcon3 detector in linear mode.

For the structure of PtdIns3K-C1 with ADP-MgF3, PtdIns3K-C1 at a final concentration of 3.6 µM was incubated on ice for 30 min with 3.6 µM NRBF2 MIT, in a buffer containing 25 mM HEPES pH 8.0, 100 mM NaCl, 1 mM TCEP pH 7.4, 4 mM CHAPSO, 0.005% (v/v) Nonidet P-40 substitute, 13 mM MgCl2, 22.5 mM NaF and 2.5 mM ADP. Grids were prepared and screened as detailed above.

### Cryo-electron microscopy data collection

For the structure of PtdIns3K-C1 with GABARAP, two datasets with identical parameters were collected on an identical sample, one with 4,776 movies and another with 13,450 movies. Micrographs were collected on a Titan Krios G1 TEM (Thermo Fisher) with a Gatan K3 detector (Gatan) in counting mode and energy filter set at 20 eV, using a 100 µm objective aperture. The magnification was 105,000x and the pixel size of 0.725 Å/px. Micrographs were collected with EPU (Thermo Fisher) in AFIS mode, with one movie of 40 frames per hole, at a defocus range –0.8 to 2 µm. The dose rate was 27 e^-^/px/s over an exposure length of 0.97 s, making 50 e^-^/Å^2^ total dose. See Table 4 for the statistics of the cryo-EM data collection and processing.

**Table 4:**
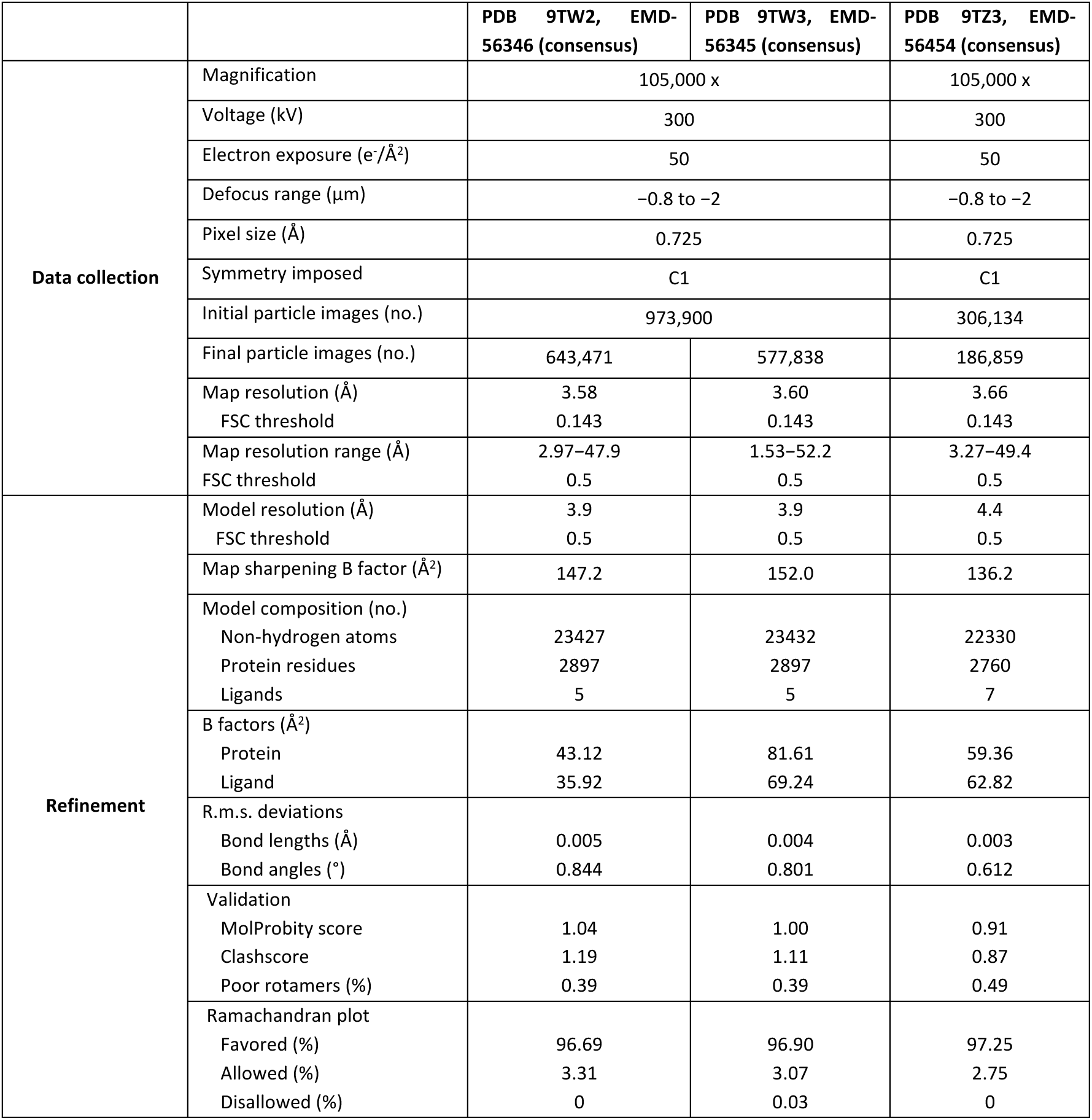
Cryo-EM data collection, refinement and validation statistics.

For the structure of PtdIns3K-C1 with ADP-MgF3, 8,483 movies were collected on a Titan Krios G1 TEM (Thermo Fisher) with a Gatan K3 detector (Gatan) in counting mode and energy filter set at 20 eV, using a 100 µm objective aperture. The magnification was 105,000x and the pixel size of 0.725 Å/px. Micrographs were collected in EPU (Thermo Fisher) in AFIS mode, with one movie of 50 frames per hole, at defocus range –0.8 to 2 µm. The dose rate was 16 e^-^/px/s over an exposure length of 1.66 s, making 50 e^-^/Å^2^ total dose. See Table 4 for the statistics of the cryo-EM data collection and processing.

### Cryo-electron microscopy data processing and model building

For the structure of PtdIns3K-C1 with GABARAP, both datasets were separately imported in CryoSPARC^59^. The movies were corrected for beam-induced motion using patch motion correction^60^ and their CTF estimated with the CTFFIND4 wrapper^61^. Movies with CTF-estimated resolution below 10 Å were selected for further processing. Particle picking was done in crYOLO-1.7.5^62^, using a custom-trained model from a previous study^37^. A total of 973,900 particles were combined in CryoSPARC and extracted at a box size of 452 px, then downsampled to 226 px. Particles were subjected to one round of 2D classification, and 932,517 particles were separated from obvious noise. Particles were aligned in a non-uniform refinement^63^, using a density of PtdIns3K-C1 and MIT as a 30 Å-low passed reference (EMD-54326, unpublished data). Four masks of different regions of the complex were created in UCSF ChimeraX^64^: 1) PIK3C3 and PIK3R4 kinases, 2) The core of PtdIns3K-C1 (PIK3R4 WD40, PIK3C3 C2 domain and helical domain), 3) The PIK3R4 helical solenoid, BECN1 and ATG14 N-termini (NTDs) and GABARAP, and 4) The BECN1 and ATG14 C-termini (CTDs). The masks were generated in CryoSPARC, and used for focused 3D classification of all particles, followed by a local refinement on the best 3D classes with unbinned particles at box size 452 px. The particles that went through a local refinement of the BECN1 and ATG14 NTDs went through an additional round of focused 3D classification and local refinement, using a smaller mask on the GABARAP protein done in UCSF ChimeraX. For the focused 3D classification of the CTDs of BECN1 and ATG14 (the adaptor arm of PtdIns3K-C1), 2 classes with a different conformation (‘main’ and ‘alternative’) were separately refined in local refinements. All particles were used in CryoSPARC non-uniform refinements to generate two consensus maps, using one class of the adaptor arm conformation or the other. Using the same particles as for the consensus maps, two composite maps were generated with TEMPy-ReFF^65^ using the five local refinements. See Table 4 for the statistics of the cryo-EM data collection and processing, and Supplementary Fig. 6 for a detailed diagram of the processing workflow. An initial model of all proteins was generated using AlphaFold3^66^. This model was fitted to each composite map by rigid-body fit in UCSF ChimeraX followed by manual adjustment assisted by local fit using the ISOLDE plugin^67^. The models were then refined using TEMPy-ReFF, followed by real space refinement in PHENIX^68^. The statistics presented in Table 4 were determined in comparison to the consensus maps using MolProbity^69^ and PHENIX, and the local resolutions and FSC resolutions determined in CryoSPARC.

For the structure of PtdIns3K-C1 with ADP-MgF3, the movies were imported in CryoSPARC, motion-corrected and CTF-estimated as above. The trained crYOLO model was used to pick 306,134 particles, which were imported in CryoSPARC and extracted at a box size of 452 px, then downsampled to 150 px. One round of 2D classification was used to get rid of obvious noise, then particles were aligned with a non-uniform refinement using the map of PtdIns3K-C1 and GABARAP (EMD-56346) obtained above as a 30 Å-low passed reference. Three masks of different regions of PtdIns3K-C1 were created in UCSF ChimeraX and CryoSPARC: 1) The core of PtdIns3K-C1 including ATG14 and BECN1 N-termini, 2) PIK3C3 and PIK3R4 kinases, and 3) ATG14 and BECN1 C-termini. The masks were used for three focused 3D classifications on all particles, followed by a local refinement on the best classes with unbinned particles at a box size of 452 px. For the PIK3C3 and PIK3R4 kinases, a non-uniform refinement was done instead of a local refinement and a smaller mask on the kinases was applied for a focused 3D classification, followed by a local refinement using the same mask. All local refinements were combined in a composite map in TEMPy-ReFF. The model of PtdIns3K-C1 (PDB 9TW2) was modified and refined as detailed above. See Table 4 for the statistics of the cryo-EM data collection and processing, and Supplementary Fig. 7 for a detailed diagram of the processing workflow.

### Hydrogen-deuterium exchange coupled to mass spectrometry

HDX experiments were performed as follows: PtdIns3K-C1 was preincubated for 30 min on ice in the absence or presence of GABARAP, and exposed for different deuteration times (3 s on ice, 3, 30, 300 and 3000 s at room temperature) to a 20 mM HEPES pD8, 150 mM NaCl, 0.5 mM TCEP deuterated buffer (D2O, Thermo Scientific, 166300100) to yield solutions that were 2 μM PtdIns3K-C1 alone or in presence of 50 μM of GABARAP in 80% D2O. Triplicated experiments were performed for each timepoint. Exchange reactions were quenched in 2 M guanidine-hydrochloride, 100 mM glycine, formic acid (Fisher Chemical, 10596814), LC-MS grade water (Romil, H949), pH 2.3 (final pH was 2.5). Samples were then flash-frozen in liquid nitrogen after quenching and stored at −80 °C until analysis. Prior to LC-MS analysis, samples were quickly thawed and injected on a HDX Manager coupled to an Acquity UPLC M-Class system (Waters) set at 0.1 °C. Samples were then digested at 15 °C using a 20 mm x 2.0 mm home-made pepsin column (Pepsin from Thermo Scientific, 20343 and hardware from Upchurch Scientific, C130-B) and loaded on a UPLC pre-column (ACQUITY UPLC BEH C18 VanGuard pre-column, 2.1 mm I.D. x 5 mm, 1.7 μM particle diameter, Waters, 186003975) for 2 min at 100 μL/min. Digested peptides were then eluted over 20 min from the pre-column onto an ACQUITY UPLC BEH C18 column (1.0 mm I.D. x 100 mm, 1.7 μM particle diameter, Waters, 186002346) using a 5−43% gradient of Acetonitrile (Romil, H050), 0.1% formic acid. MS data were acquired using a Synapt G2Si-HDMS (Waters) with electrospray ionization, using the data-independent acquisition mode (MS^E^) over a *m*/*z* range of 50–2000 and a 100 fmole/μL Glu-FibrinoPeptide solution was used for lock-mass correction and calibration. The following parameters were used during the acquisition: capillary voltage, 3 kV; sampling cone voltage, 40 V; source temperature, 90°C; desolvation gas, 150°C and 650 L.h^−1^; scan time, 0.3 s; trap collision energy ramp, 20–45 eV. Peptide identification was performed using ProteinLynx Global Server 3.0.1 (PLGS, Waters) with a home-made protein sequence library containing PIK3C3, PIK3R4, ATG14, BECN1, GABARAP, and pepsin sequences, with peptide and fragment tolerances set automatically by PLGS. Peptides were then filtered using HDExaminer 3.4.0 (Trajan Scientific and Medical) with a minimum fragment per amino acid of 0.2, a minimum intensity of 10^3^, a peptide length between 5 and 25 residues, a minimum PLGS score of 6.62, and an MH^+^ tolerance of 5 ppm. An initial automated spectral processing step was conducted by HDExaminer followed by a manual inspection of individual peptides for sufficient quality where only one charge state per peptide was kept. Deuterium uptakes were not corrected for back-exchange and are reported as relative. HDX-MS results were statistically validated using an in-house program (Archaeopteryx), where significant differences required 0.3 Da, 5 %, and *p*-value < 0.05 (*n*=3) using an unpaired two-tailed *t*-test. HDX-MS results were illustrated on the PDB 9TW2 model using PyMOL 2.5.4 (Schrödinger). Deuteration tables are available in Supplementary Data 1. The mass spectrometry proteomics data (including raw data, tables of every peptide included within the dataset and all relative uptake plots) have been deposited to the ProteomeXchange Consortium via the PRIDE partner repository^70^ with the dataset identifier PXD074963.

### Large unilamellar vesicles (LUVs) preparation

The lipid stocks of Table 2 were mixed to prepare the lipid mixture ADluv15 at 5 mg/mL in chloroform, as shown in Table 3. The solvent was then evaporated with a stream of nitrogen gas and desiccated for 1 h under vacuum. The dried lipid film was resuspended in a buffer with 25 mM HEPES pH 7.0, 150 mM NaCl for a final concentration of 5 mg/mL. The liposomes were vortexed for 2 min (Vortex Genie 2, Scientific Industries), followed by a sonication for 1 min in a water bath sonicator (Ultrawave U50). Liposomes were frozen and thawed 10 times in a cycle between liquid nitrogen and a 43°C water bath, followed by an extrusion for 30 cycles using a 100 nm diameter extruder (Whatman Anotop 10 syringe filter 0.1 μm pore size, 68091112). Liposomes were aliquoted, flash-frozen in liquid nitrogen and stored at –80°C.

### In vitro reconstitution of the mATG8 lipidation on LUVs

For the maleimide lipidation in the XL-MS experiment, 0.5 mg/mL of ADluv15 were incubated with 10 µM of GABARAP (L117C) in a buffer containing 25 mM HEPES pH 7.0 and 150 mM NaCl at 4°C overnight. Next day, the reaction was quenched with 5 mM 2-mercaptoethanol (Aldrich M6250) and the LUVs aggregates removed by pelleting with a centrifugation at 1,000 g for 1 min at 4°C. The LUVs in the supernatant were concentrated by a 20 min centrifugation at 21,000 g at 4°C, and resuspended in 1/10 of the initial reaction volume.

### Crosslinking Mass Spectrometry

For the sample with soluble GABARAP, 4 µM of PtdIns3K-C1 were incubated with 50 µM of GABARAP (1−116) for 30 min on ice in a buffer containing 20 mM HEPES pH8, 150 mM NaCl and 0.5 mM TCEP. For the sample containing LUVs, the vesicles were coated with GABARAP (L117C) by a maleimide reaction as detailed above. PtdIns3K-C1 was then added to the concentrated LUVs at a concentration of 3.6 µM, for 50 µL final volume, and incubated on ice for 30 min. Protein crosslinking reactions were carried out by adding 2 mM of sulfo-SDA to the protein mixture, with 5 min incubation on ice, followed by 10 s 365 nm UV radiation from a home build UV LED setup. Crosslinking reactions were quenched with the addition of ammonium bicarbonate to a final concentration of 50 mM. The quenched solution was reduced with 5 mM DTT and alkylated with 20 mM idoacetamide. SP3 protocol71,72 was used to clean-up and buffer exchange the reduced and alkylated protein, shortly, proteins were washed with ethanol using magnetic beads for protein capture and binding. The proteins were resuspended in 100 mM NH4HCO3 and were digested with trypsin (Promega) at an enzyme-to-substrate ratio of 1:20, and protease max 0.1% (Promega). Digestion was carried out overnight at 37 °C. Clean-up of peptide digests was carried out with HyperSep SpinTip P-20 (ThermoScientific) C18 columns, using 60 % Acetonitrile as the elution solvent. Peptides were then evaporated to dryness *via* Speed Vac Plus (Savant). Dried peptides were resupended in 30 % acetonitrile and were fractionated *via* size exclusion chromatography using a Superdex 30 Increase 3.2/300 column (GE heathcare) at a flow rate of 20 µL/min using 30% (v/v) ACN 0.1 % (v/v) TFA as a mobile phase. Fractions were taken every 5 min, and the 2nd to 7th fractions containing crosslinked peptides were collected. Dried peptides were suspended in 3% (v/v) Acetonitrile and 0.1 % (v/v) formic acid and analysed by nano-scale capillary LC-MS/MS using an Ultimate U3000 HPLC (ThermoScientific) to deliver a flow of 300 nL/min. Peptides were trapped on a C18 Acclaim PepMap100 5 μm, 0.3 μm x 5 mm cartridge (ThermoScientific) before separation on Aurora Ultimate C18, 1.7 μm, 75 μm x 25 cm (Ionopticks). Peptides were eluted on optimized gradients of 90 min and interfaced *via* an EasySpray ionization source to a tribrid quadrupole Orbitrap mass spectrometer (Orbitrap Eclipse, ThermoScientific) equipped with FAIMS. MS data were acquired in data dependent mode with a Top-25 method, high resolution scans full mass scans were carried out (R=120,000, *m/z* 400–1550) followed by higher energy collision dissociation with stepped collision energy range 21, 30, 34 % normalized collision energy. The tandem mass spectra were recorded (R=60,000, isolation window *m/z* 1, dynamic exclusion 50 s). Mass spectrometry measurements were cycled for 3 s durations between FAIMS CV –45, and –60 V. Cross linking data analysis was done by converting Xcalibur raw files to MGF files using ProteoWizard73 and crosslinks were analysed by XiSearch74. Search conditions used 3 maximum missed cleavages with a minimum peptide length of 5. Variable modifications used were carbmidomethylation of cysteine (57.02146 Da) and Methionine oxidation (15.99491 Da). False discovery rate was set to 5%. Data was imported in XiView75 for manual filtering, and exported to the PDB 9TW2 in PyMOL 2.5.4 (Schrödinger). The mass spectrometry proteomics data have been deposited to the ProteomeXchange Consortium via the PRIDE partner repository with the dataset identifier PXD074924.

### WIPI2 and FIP200 colocalization experiments

HEK293 cells were plated on glass coverslips to be 60–80% the next day for the experiment. Treatments with PP242 and the ATG7 inhibitor were done by withdrawing half the volume of the medium in each well, adding the compound at 2x final concentration and then adding the solution back to the well for the indicated times. After fixation in 3.7 % formaldehyde the coverslips were permeabilised in 0.1 % NP40 and stained with antibodies to endogenous LC3 (Caltag Medsystems MBL-PM036), FIP200 (ProteinTech 10069-1-AP) and WIPI2 (Bio Rad LaboratoriesMCA5780GA) as indicated. Images were acquired on a ZEISS Axio Imager D2 and analysed using ImageJ.

### Multiple sequence alignment

mATG8 proteins sequences were obtained from UniProt (https://www.uniprot.org/)76 and aligned using Clustal Omega (https://www.ebi.ac.uk/jdispatcher/msa/clustalo)77. Both the multiple sequence alignment and the phylogenic tree of Supplementary Fig. 5 were retrieved from Clustal Omega.

## Data availability

Plasmids used in this study (Table 1) were deposited on Addgene (https://www.addgene.org/). All the cryo-EM maps have been deposited on the Electron Microscopy Data Bank (EMDB). The deposition codes for the GABARAP dataset are the following: EMD-56357 (composite map), EMD-56346 (consensus map), EMD-56347, EMD-56348, EMD-56350, EMD-56351, EMD-56353 for the main conformation; and EMD-56358 (composite map), EMD-56345 (consensus map) and EMD-56343 for the adaptor arm alternative conformation (see Supplementary Fig. 6). The two atomic models were deposited on the Protein Data Bank (PDB) under the accession codes 9TW2 (main conformation) and 9TW3 (alternative conformation). For the ADP-containing dataset, the accession codes are the following: EMD-56458 (composite map), EMD-56454 (consensus map), EMD-56455, EMD-56456, EMD-56457 (see Supplementary Fig. 7). The atomic model was deposited on the PDB with accession code 9TZ3. HDX-MS and XL-MS data have been deposited to the ProteomeXchange Consortium via the PRIDE partner repository with the respective identifiers PXD074963 and PXD074924. Deuteration tables are available in Supplementary Data 1.

---

**Supplementary Figure 1.**
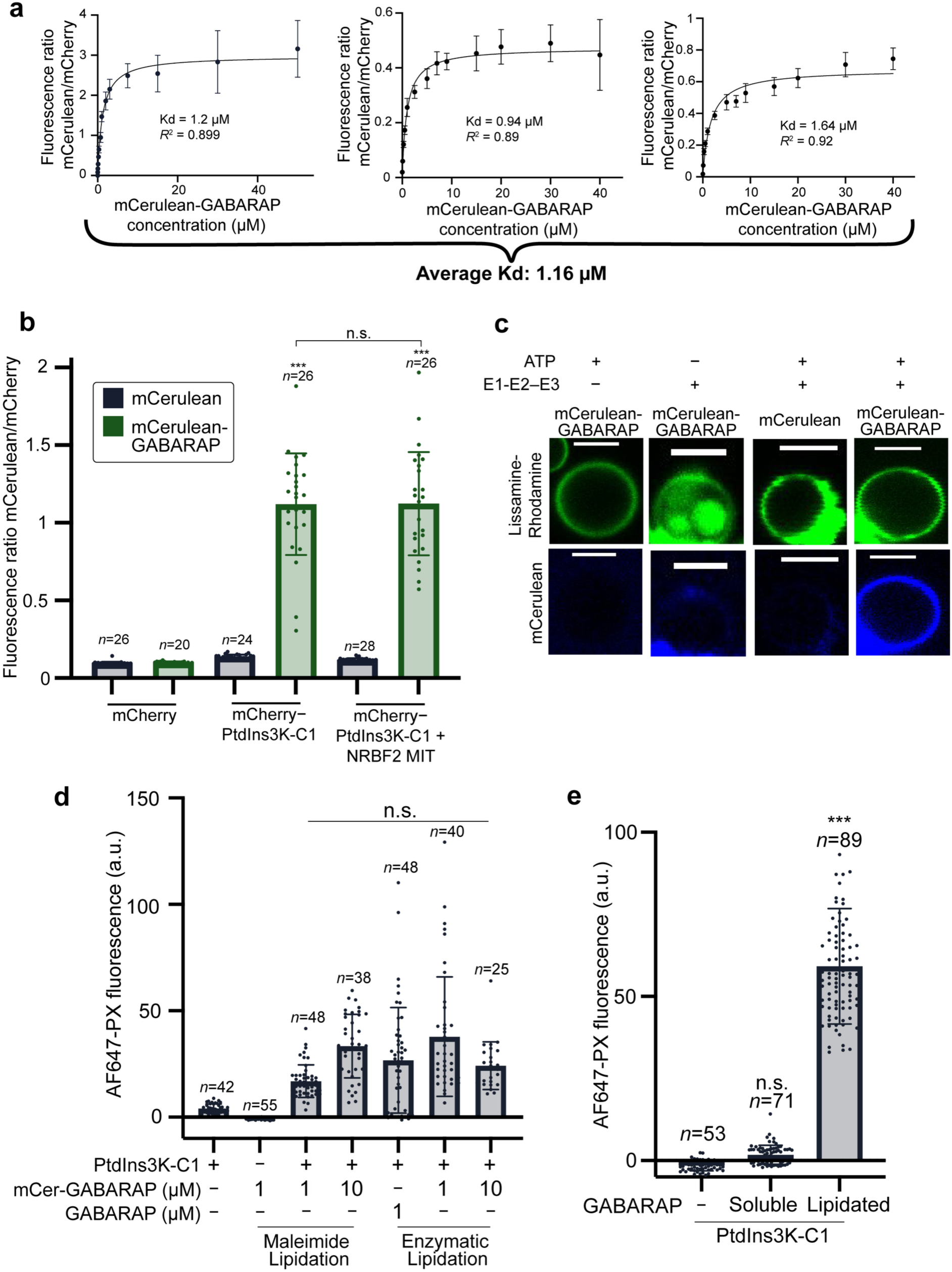
Controls for the binding and the activation of PtdIns3K-C1 with GABARAP. **a** Affinity of PtdIns3K-C1 for GABARAP. Replicates for the experiment shown in Fig. 2c. The plots show the average of at least 29 beads ± *SD*. A Kd and associated *R*2 were determined by a non-linear fit. **b** mCherry, mCerulean and NRBF2 MIT do not affect the binding between PtdIns3K-C1 and GABARAP. mCherry or mCherry−PtdIns3K-C1 were immobilized to beads as presented in Fig. 2a, and incubated with mCerulean or mCerulean-GABARAP, in the presence or the absence of the NRBF2 MIT domain. **c** The lipidation of mCerulean-GABARAP to GUVs is not caused by the mCerulean protein, and needs ATP and the lipidation enzymes E1-E2−E3. Representative confocal micrographs of GUVs, in the Lissamine-Rhodamine (lipids) and mCerulean channels. Scale bars: 5 µm. **d** Comparison of the enzymatic and maleimide-based GABARAP lipidations for PtdIns3K-C1 activity. mCerulean (mCer)-GABARAP was conjugated to GUVs through maleimide reaction or the enzymatic lipidation cascade, then incubated with PtdIns3K-C1 and AF647-PX to follow PtdIns3P production in an endpoint. A control with GABARAP instead of mCer-GABARAP shows the mCerulean does not affect PtdIns3K-C1 activation. Average of the AF647-PX fluorescence intensity in arbitrary units (a.u.) for *n* GUVs ± *SD.* Averages were compared to the sample with PtdIns3K-C1 but without GABARAP with a one-way ANOVA with Tukey’s correction for multitesting (n.s.: *p* > 0.05, *: *p* < 0.05, ***: *p* < 0.0001). **e** Soluble GABARAP does not activate PtdIns3K-C1. GUV-based endpoint PtdIns3K-C1 activity assay using GUVs with soluble or maleimide-lipidated GABARAP. Average of the AF647-PX fluorescence intensity for *n* GUVs ± *SD.* Means were compared to the PtdIns3K-C1 only control with a one-way ANOVA with Tukey’s correction for multitesting (n.s.: *p* > 0.05, ***: *p* < 0.0001).

**Supplementary Figure 2.**
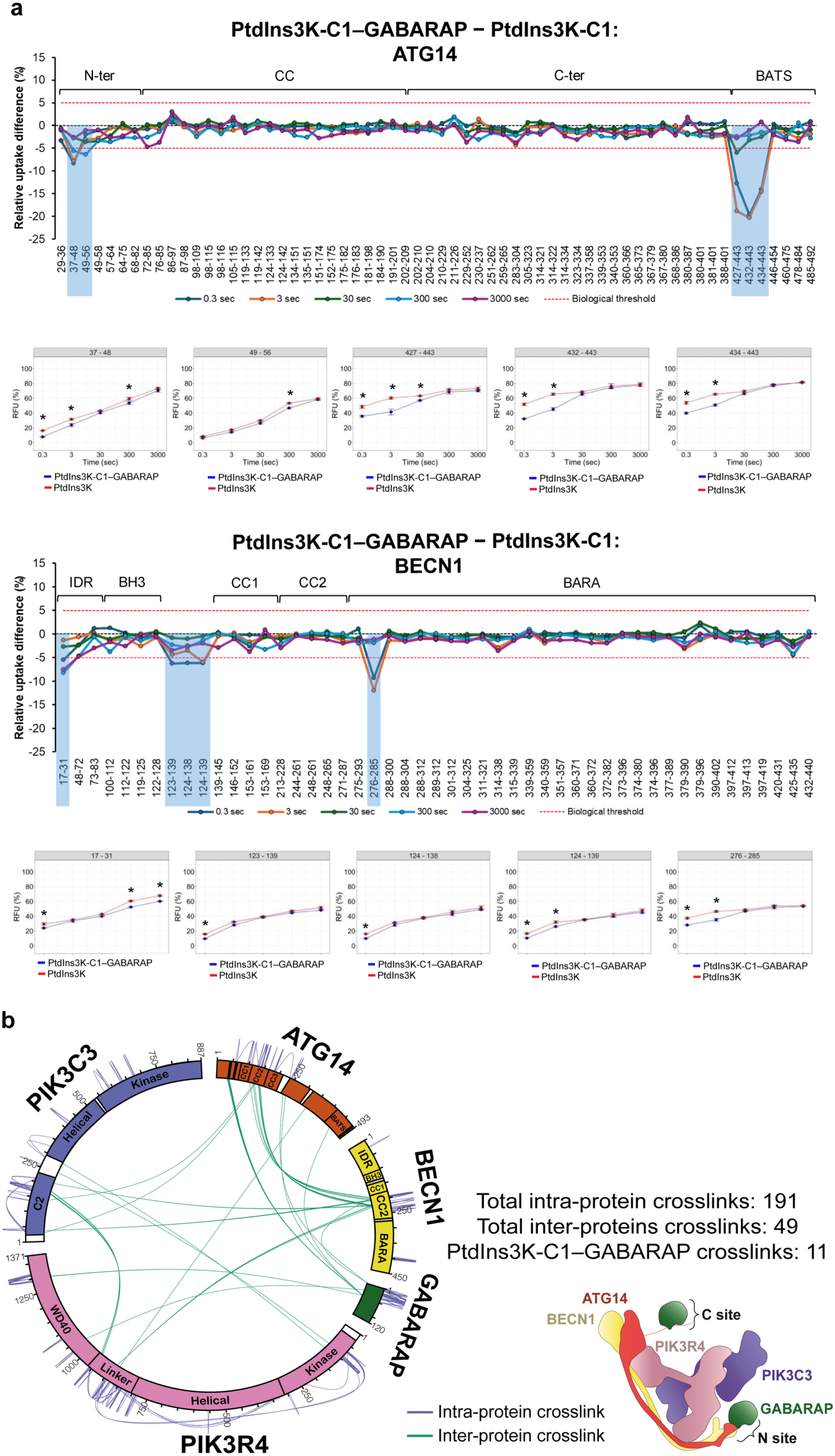
Identification of two GABARAP-binding sites on PtdIns3K-C1 by a combination of structural approaches. **a** HDX-MS difference plots for the ATG14 and BECN1 subunits. Peptides showing significant differences (PtdIns3K-C1−GABRAP – PtdIns3K-C1) are framed in blue (protection) and their corresponding uptake plots are displayed. Stars indicate timepoints showing significant differences. No significant differences have been identified for the PIK3R4 and PIK3C3 subunits. **b** Crosslinking-MS maps on PtdIns3K-C1 subunits. Identified crosslinked peptides between PtdIns3K-C1 and soluble GABARAP are mapped on the different protein domains. Numbers of intra-protein crosslinks (blue) and inter-protein crosslinks (green) are indicated.

**Supplementary Figure 3.**
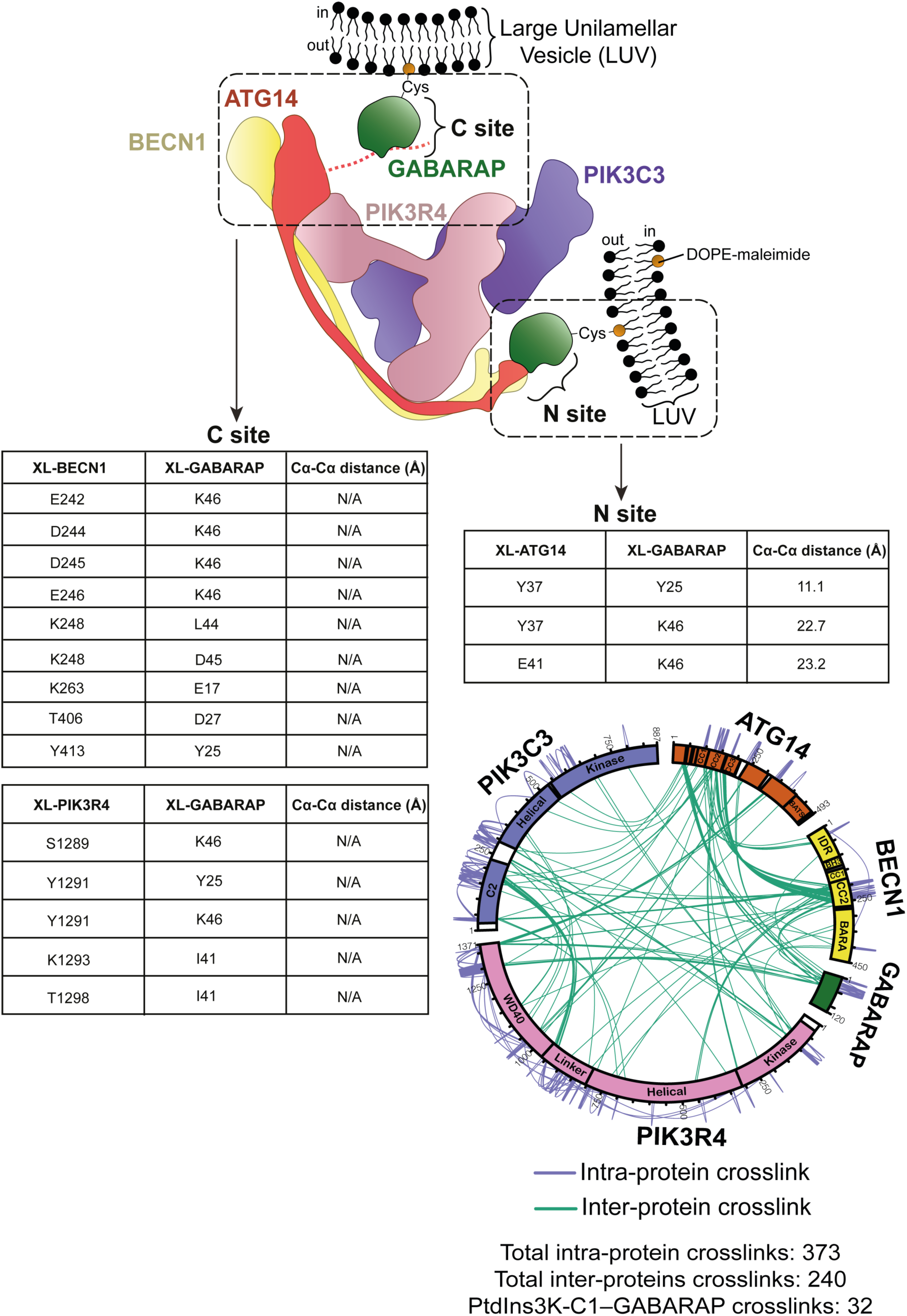
PtdIns3K-C1 binds GABARAP in the C site and the N site on membranes. Inter-crosslinked peptides between PtdIns3K-C1 and lipidated GABARAP have been identified in both the N site (to ATG14) and the C site (to PIK3R4 and BECN1) in the presence of 100 nm Large Unilamellar Vesicles (LUVs). GABARAP was covalently coupled to the LUVs by maleimide coupling (Cys = cysteine). LUVs inner and outer membrane leaflets are labelled as ‘in’ and ‘out’, respectively. All identified crosslinked peptides between PtdIns3K-C1 and GABARAP are mapped on the different protein domains. Numbers of intra-protein crosslinks (blue) and inter-protein crosslinks (green) are indicated.

**Supplementary Figure 4.**
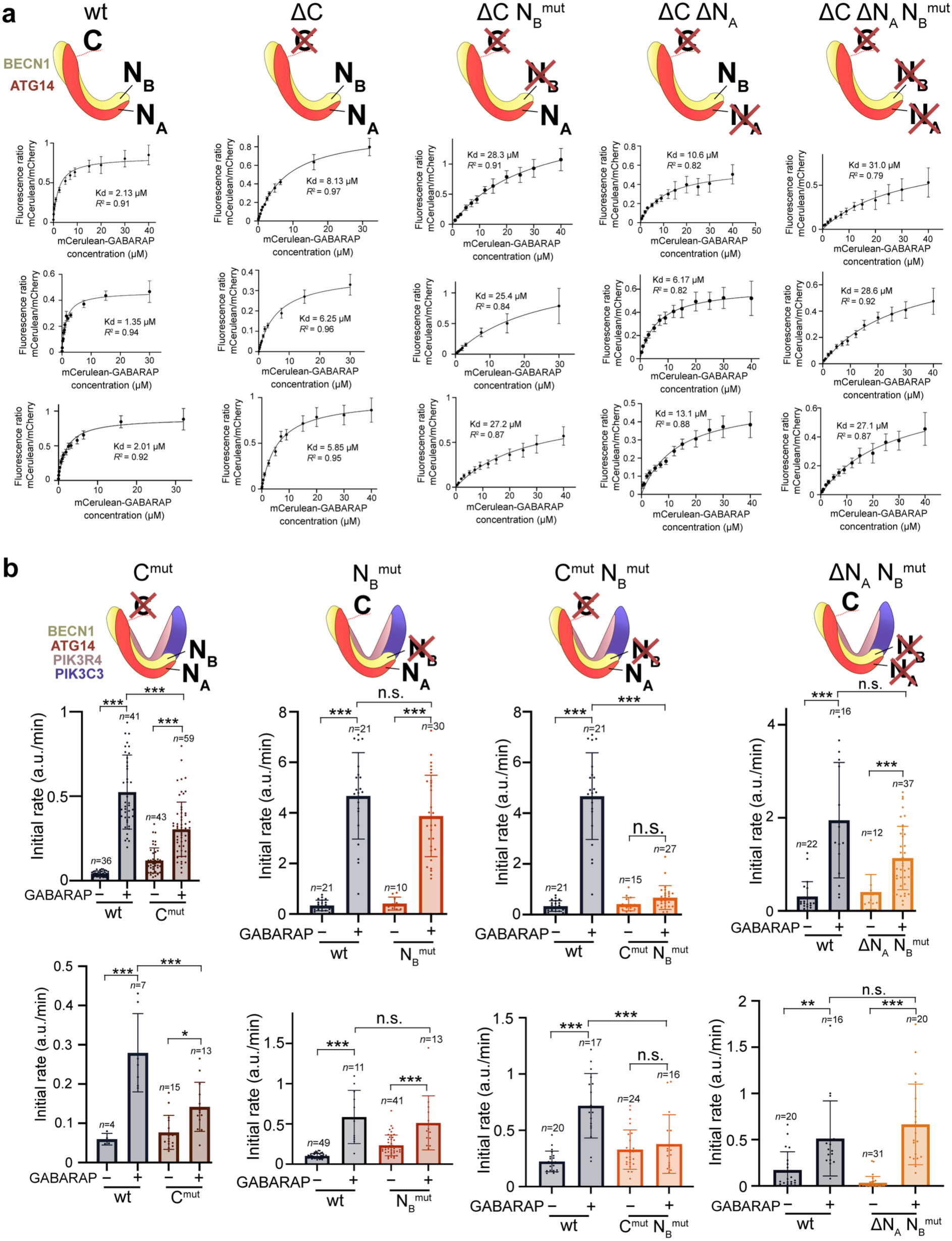
The C and N sites cooperate for PtdIns3K-C1 activation by GABARAP. **a** Replicates of affinity for GABARAP of different C site and N site mutants as shown in Fig. 6b. mCherry-ATG14−BECN1 constructs were immobilized on anti-RFP beads and incubated with different concentrations of mCerulean-GABARAP. Kd and R2 were determined by a non-linear regression fit. **b** Replicates of GUV-based activity assay of PtdIns3K-C1 carrying mutations on C and N sites in presence or absence of maleimide-lipidated GABARAP, two additional replicates of the experiments shown in Fig. 6c. PtdIns3P production initial rates expressed in AF647-PX fluorescence/min, average ± *SD* for *n* GUVs. Means were compared in a two-way ANOVA with Tukey’s correction for multitesting (ns: *p* > 0.05, *: *p* < 0.05, **: *p* < 0.002, ***: *p* < 0.0001).

**Supplementary Figure 5.**
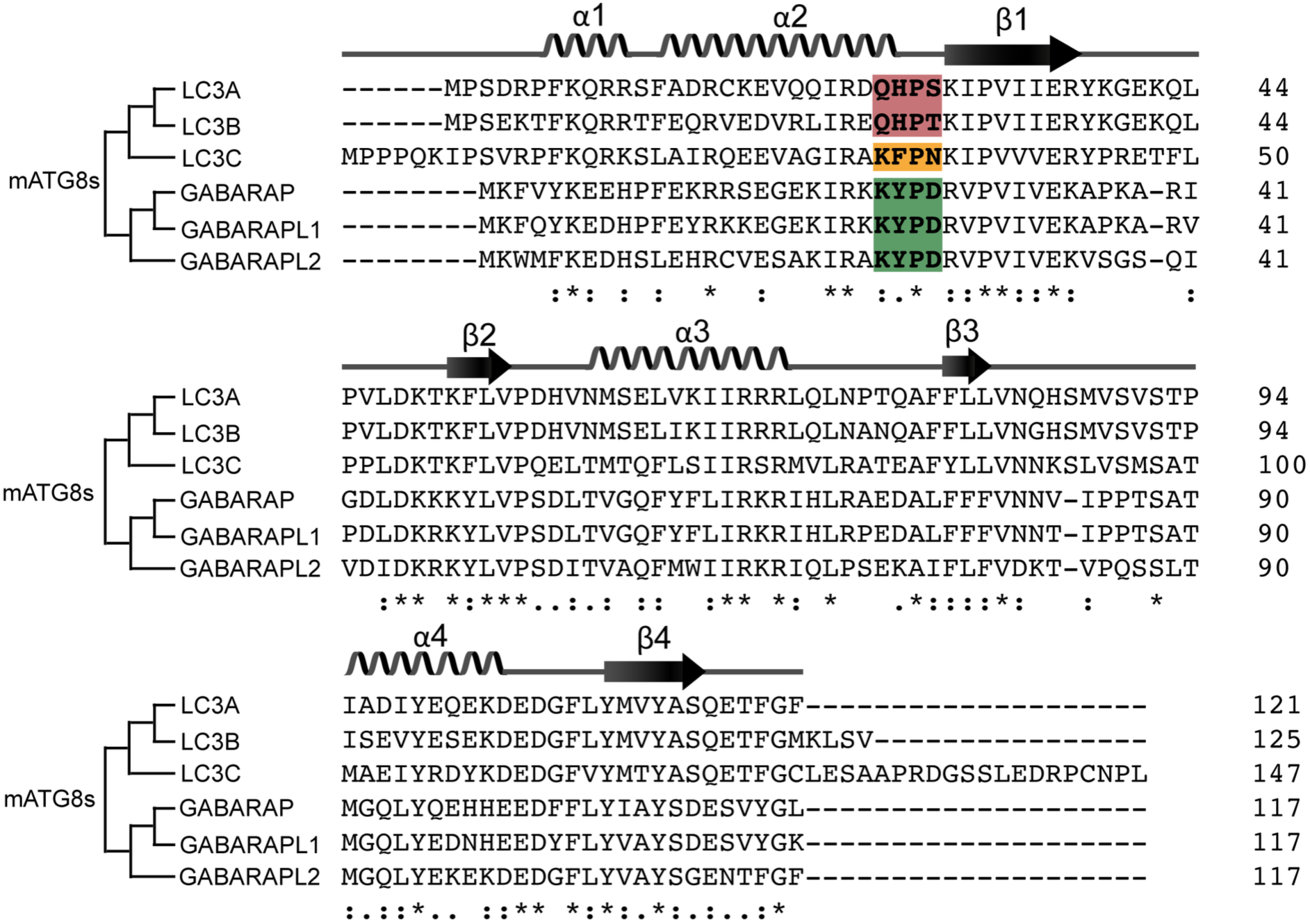
LC3 and GABARAP families differ in the ɑ2-β1 loop binding PtdIns3K-C1 NA site. Multiple sequence alignment using Clustal Omega is overlaid with the secondary structures of GABARAP (PDB 1GNU) and the phylogenic tree of all mATG8s. The KYPD sequence found in GABARAP family is highlighted in green, the similar sequence KFPN of LC3C highlighted in orange, and the QHPS/T sequence of LC3A/B highlighted in red. ‘`*`’: fully conserved residues, ‘:’: conservation with strongly similar properties, ‘.’: conservation with weakly similar properties, ‘ ‘: non-conserved residues.

**Supplementary Figure 6.**
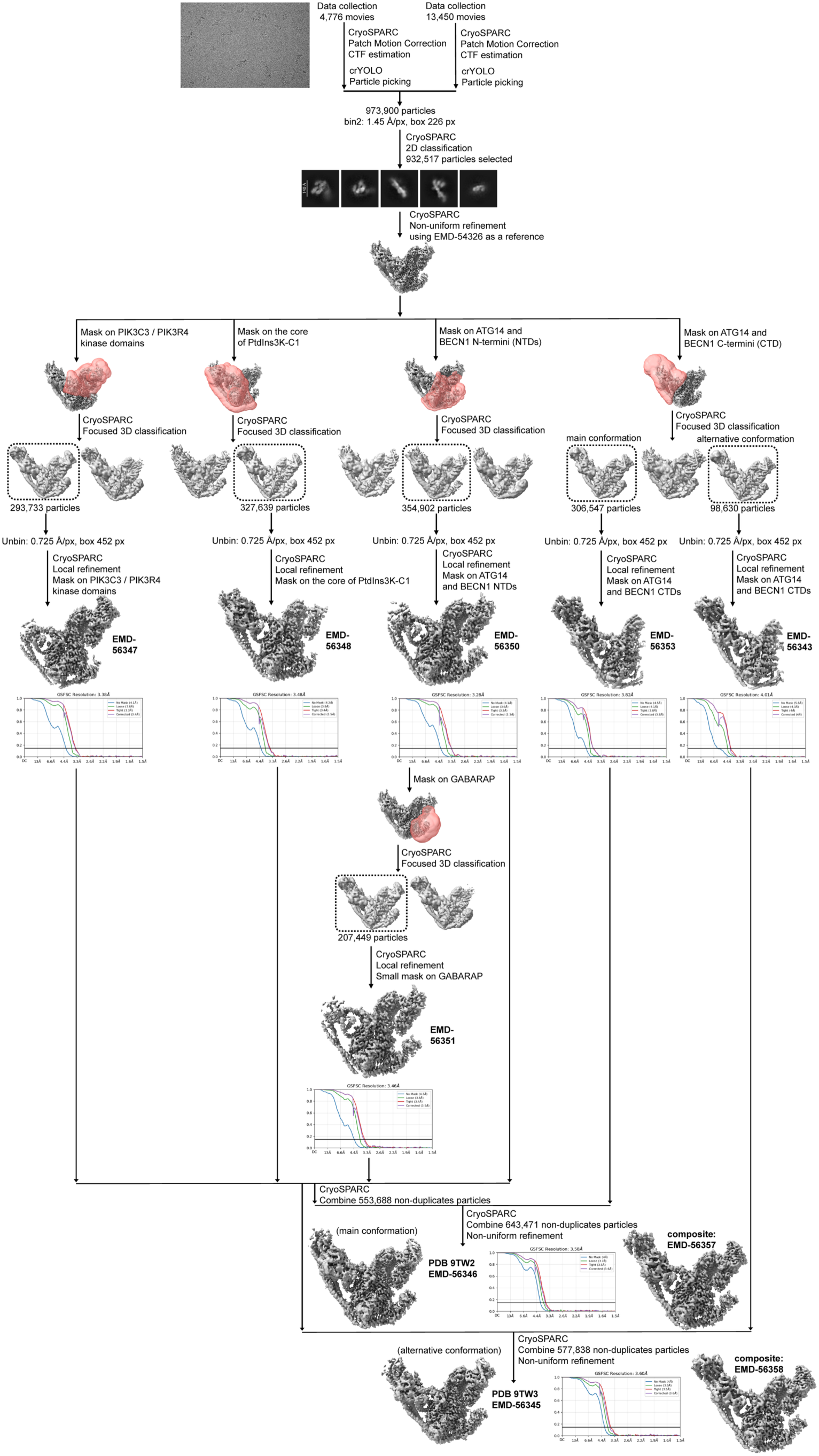
Single particle cryo-EM processing workflow for the structure of PtdIns3K-C1 with NRBF2 MIT and GABARAP. Only a few representative 2D and 3D classes are shown. EMDB accession codes are written under their corresponding maps, as well as the PDB code of the fitted model.

**Supplementary Figure 7.**
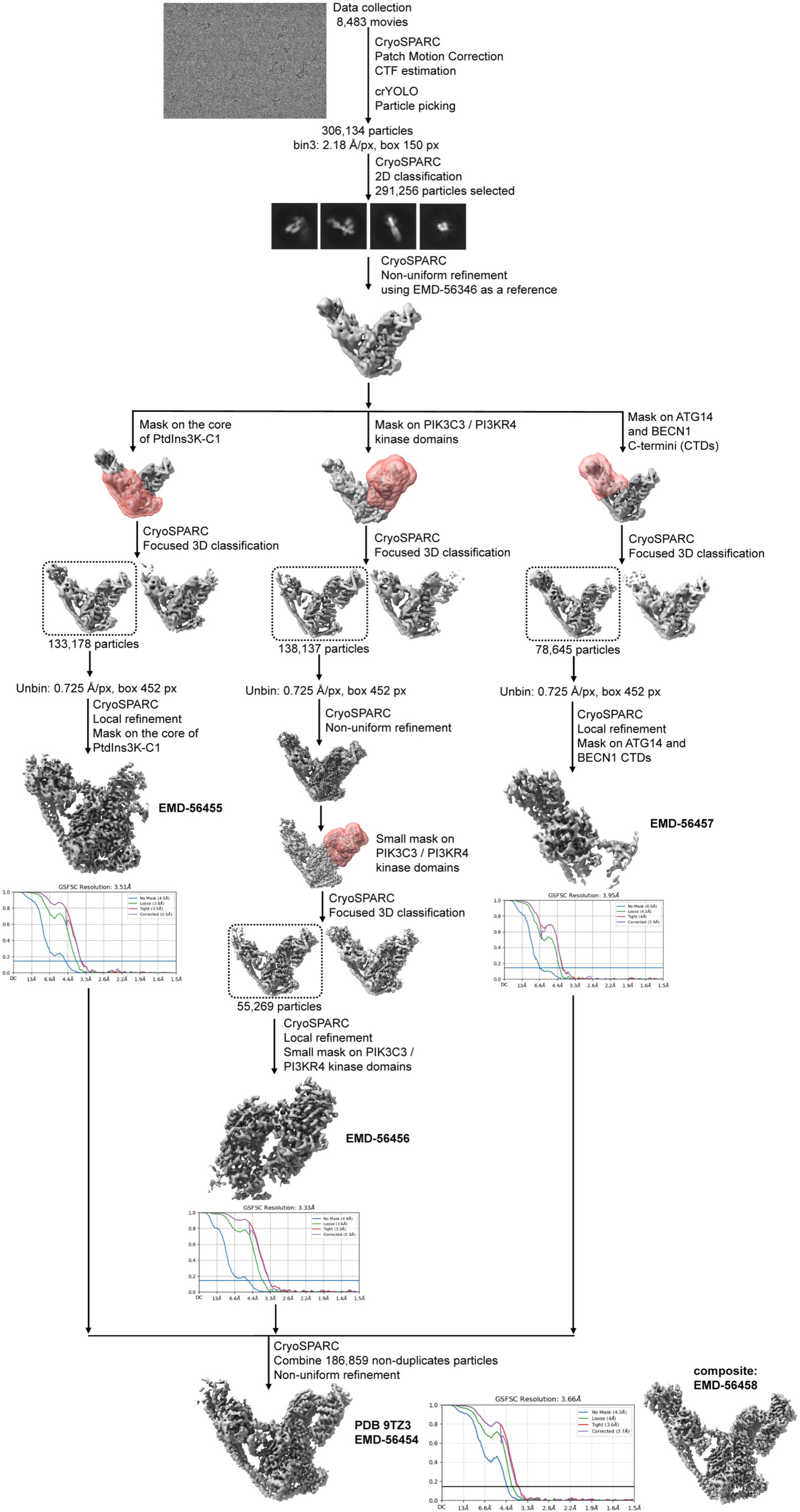
Single particle cryo-EM processing workflow for the structure of PtdIns3K-C1 with NRBF2 MIT and ADP-MgF3. Only a few representative 2D and 3D classes are shown. EMDB accession codes are written under their corresponding maps, as well as the PDB code of the fitted model.

